# Multiscale mechanisms driving tissue rupture by invading cells

**DOI:** 10.1101/2023.09.25.559247

**Authors:** Selwin K. Wu, Fuqiang Sun, Celestine Z. Ho, Yuting Lou, Christina Bao-Xian Huang, Mui Hoon Nai, Jingwei Xiao, Murat Shagirov, Jasmine Fei Li Chin, Diana Lim, Suzie Verma, David SP Tan, Philippe Marcq, Alpha S. Yap, Chwee Teck Lim, Tetsuya Hiraiwa, Yuan Lin, Boon Chuan Low

**Author notes:** Contributed Equally (Selwin K. Wu, Fuqiang Sun, Celestine Z. Ho, Yuting Lou). Correspondence & &.

## Abstract

Cells migrate and invade tissues during development, immune responses, and cancer. Collective invasion is generally understood to be driven by invading cells unjamming and pushing through barriers such as the extracellular matrix and surrounding tissues. Whether these barriers actively contribute to invasion remains unclear. Using ovarian adenocarcinoma spheroids invading mesothelium derived from benign pleural effusions as an experimental model, combined with modelling, we examine invasion across molecular to multicellular scales. We identify intercellular integrin adhesions linking invasive leader cells to the tissue barrier, triggering apical constrictions within the barrier. This constriction shrinks cell-cell contacts, leading to barrier rupture. Thus, the tissue barrier plays a mechanically active role in invasion. Rather than cells pushing through, we find that coordinated subcellular contractility between the invading leader cell and the barrier drives barrier tensile rupture and invasion, independent of a jamming transition. Together, our findings challenge prevailing paradigms of collective cell invasion.

## Introduction

Cell migration and invasion into tissues and the extracellular matrix are fundamental for the development and maintenance of organisms and are rampant in diseases such as cancer^1,2^. During cancer, tumour cells traverse into foreign tissues during metastasis^2^, while in development, undifferentiated cells move into tissues to form different tissue types^3^.

A variety of microenvironments contain the extracellular matrix and closely packed cells adherent to one another^4,5^. Whether these mechanical barriers actively contribute to mammalian tissue invasion and whether cell-matrix adhesion receptors, such as integrins, are involved remains unclear. Since tumour cell clusters are more capable of forming secondary metastases than single cells^6^. Another key challenge is to identify the changes in subcellular dynamics that link molecular mechanisms to cellular and multicellular rearrangements in collective invasion of tissues.

Our current understanding of how cells collectively invade tissues stems from the study of angiogenesis, neural crest migration and border cell migration. Landmark studies in Drosophila showed that the microtopography of barrier tissue provides an energetically favourable path of least resistance for border cell cluster migration, and cell-cell adhesion contributes to the traction of border cells^7^. Notably, mechanical feedback by E-cadherin-based Rac signalling promotes direction sensing of border cells through nurse cells^8^. Despite these insights, a deep understanding of collective invasion into mammalian tissues is lacking. An interdisciplinary approach to achieve a multiscale understanding^9,10^ of collective tissue invasion in cancer remains largely unexplored.

We use three-dimensional co-cultures mimicking ovarian tumour invasion of the mesothelium as a model system in our experiments. Previously studies suggested that integrin and talin-dependent activation of non-muscle myosin-II was required for spheroids to displace the mesothelial cells apart during invasion^11,12^. Here, we study the interplay between fracture mechanics and cell invasion across molecular to multicellular scales. First, we have identified a material-based phenomenon of tissue rupture by collective invasion and the patterns of motion driving invasion. Then, we worked out the cellular and molecular mechanisms underlying the patterns of motion driving invasion and the role of multicellular dynamics during spheroid invasion. Finally, our theoretical modelling predicts the quantitative basis of the invasion process. We conclude that instead of the widely accepted model of invading cells pushing through the tissue barrier or jamming transition-mediated cell invasion, a coordinated patterning of subcellular contractility by invading cells and the tissue barrier drives tissue invasion. Thus, our findings challenge the existing paradigm of how cells invade tissues.

## Results

We study how cells migrate and invade the ECM and tissue barrier through experiments and theory. Our experiments used three-dimensional co-cultures that mimic ovarian tumour invasion of the mesothelium^11-14^. The mesothelium is a monolayer of simple squamous epithelium^15^ covering the human peritoneum’s stroma (Fig. 1a). In peritoneal metastasis^16^, the mesothelium is cleared by ovarian tumours, eventually leading to the absence of the mesothelium underneath the tumour and indentation of the stroma (Fig. 1b,b’).

**Figure 1.**
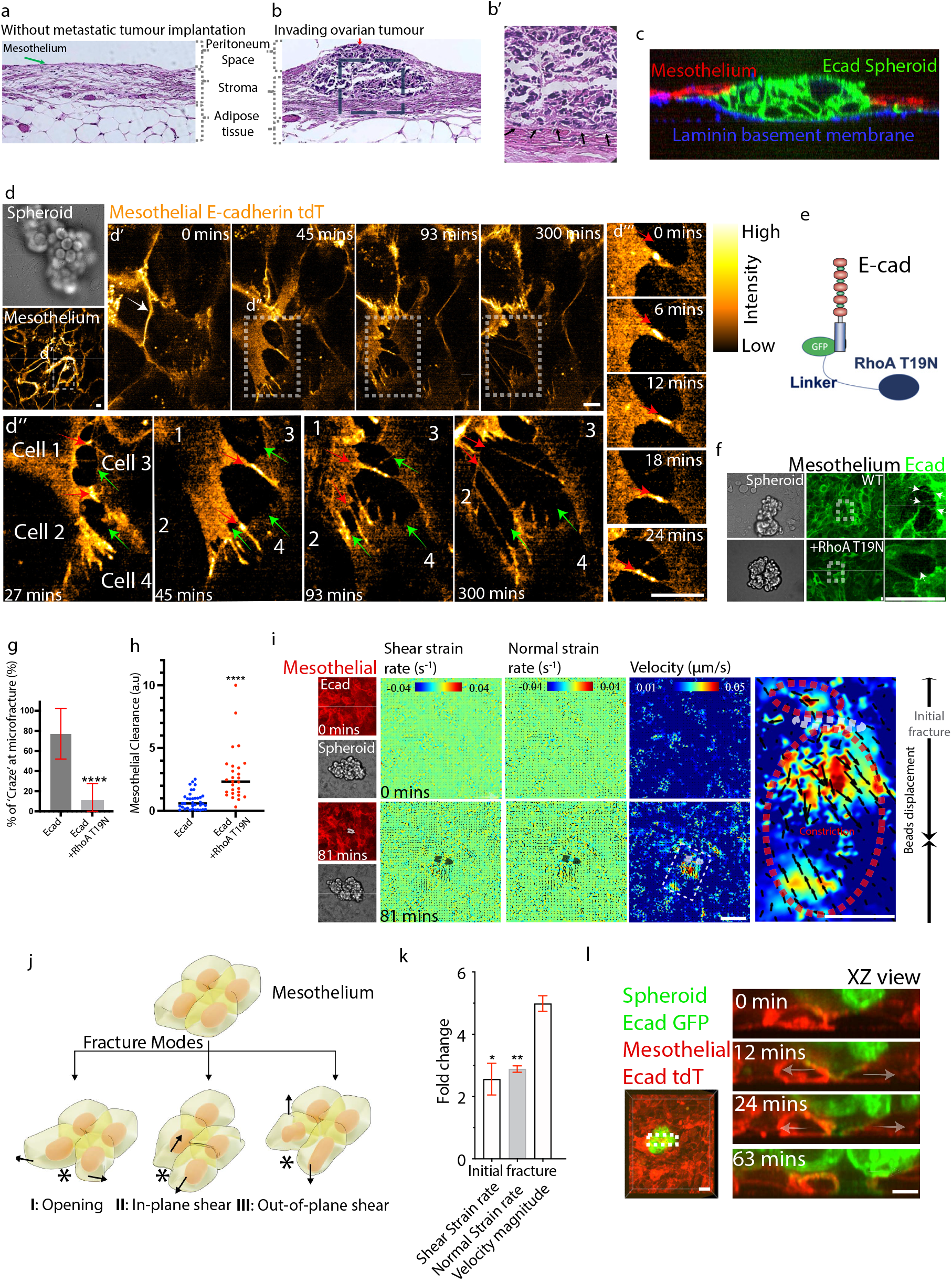
Spheroid-induced mesothelial tensile fracture. (a,b) Ovarian tumour metastasis. Hematoxylin and eosin-stained human peritoneal surface without metastatic tumour implantation (a) and a microscopic tumour implant on the omental surface from a patient with metastatic serous ovarian carcinoma (b). Mesothelium (green arrow, a) lines the abdominal surface but is not detected beneath the invading ovarian tumour (b & b’). Representative images from 4 ovarian cancer patients with different layers of tissue are indicated. Inset from b’ with arrows indicating the invading tumour/stroma interface without the mesothelium. (c) An E-cadherin GFP-expressing spheroid invades and clears the mesothelium with a basement membrane. (d) Mesothelial cell-cell contact forms E-cadherin fibrillar bridges and microvoids upon spheroid invasion. Representative time-lapse images of the mesothelium under a spheroid (colour-coded in hot orange). d’-d’’’ with detailed views. d’’’ shows a cadherin fibrillar bridge being extended as crazing proceeds. Red arrows point towards a fibrillar extension, and green arrows point towards a microvoid. (e) Schematic of E-cad-GFP-RhoA-T19N fusion protein. (f,g) Reduction of E-cad fibrillar bridges during mesothelial fracture when E-cad-based contractility is reduced. Representative images with arrows indicating fibrillar bridges (f) and quantification (g) of the percentage of mesothelium E-cad fibrillar bridges when either RhoA-T19N mutant is fused to E-cad or in controls. (h) Increased size of mesothelial clearance when E-cadherin adhesion is compromised. Quantitative comparison of the mesothelial clearance size when either RhoA-T19N mutant is fused to mesothelial E-cad or in controls. (i) Tension-induced fracture. PIV analysis of fluorescent microbeads on top of the mesothelium expressing E-cad tdTomato. The heatmaps of shear strain rate (s^-1^) or normal strain rate (s^-1^) or velocity(µm/s) from the PIV analysis. Right magnified view (white dotted rectangle) of the velocity PIV heatmap shows outward vectors (red dotted circles) away from the fracture initiation site (grey dotted circle). (j) Diagram of three different modes of fracture which the mesothelium may undergo (modified from modes of material fracture). The modes are Mode I: Opening (stress orthogonal to the local plane of the crack surface), Mode II: In-plane-shear (stress parallel to the crack surface but orthogonal to the crack front), and Mode III: Out-of-plane shear (stress parallel to the crack surface and the crack front). The arrows indicate the direction of stress, and asterisks indicate the crack point. (k) Tension-induced fracture is the dominant initiating mode of fracture. Comparison between the fold changes of the shear strain rate and normal strain rate with velocity magnitude analysis during fracture initiation and fracture. (l) Mode III: Out-of-plane shear fracture is not detected when the spheroid begins to invade the mesothelium. XZ views of time-lapse lattice light-sheet microscopy images of E-cad GFP spheroid invading E-cad tdT mesothelium. Grey arrows indicate the opening of mesothelial cell-cell contacts. All data are means ± SEM from at least n=2 independent experiments, except for (g), which are from n=8 independent experiments, ns, not significant, *p < 0.05, **p < 0.01, ****p < 0.0001; analysed with unpaired t-test. Scale bars, 5µm.

Our modified 3D culture systems can phenocopy our clinical observations (Fig. 1c). Human ovarian adenocarcinoma OVCA433 epithelial cells were cultured to form spheroids in suspension, bearing a grape cluster-like morphology (Fig. 1d)^17^. These spheroids were introduced onto the ZT mesothelia^13^ that form a laminin-rich basement membrane when grown on Matrigel (Fig. 1c). The E-cad GFP expressing spheroid invaded and cleared the underlying mesothelium (Fig. 1c), mimicking our clinical observations. Additionally, multicellular assemblies migrating into tissues can also occur as a general morphogenetic process that is not just specific to tumour invasion. This is because spheroids formed from mesothelial cells can also migrate into mesothelium grown on Matrigel (Supplementary Fig. 1a).

### Rupture initiation of the mesothelium during spheroid invasion

We first study tissue invasion at high resolution. As spheroids invade the mesothelium, a phenomenon resembling crazing is observed at mesothelial cell-cell junctions (Fig. 1d’ and Video 1). Crazing is a unique feature when thermoplastic polymer materials rupture^18^, characterized by a network of fine cracks within the material. It occurs when excessive tensile stress or stretching is applied to a polymer, producing gaps termed microvoids between fibrillar bridges that are thought to help sustain some stress (Supplementary Fig. 1b).

At the mesothelium, we observed the pulling of cell-cell junctions leading to the elongation of fibrillar bridges linked by the intercellular adhesion molecule, E-cadherin, and the formation of microvoids before mesothelial rupture (Fig. 1d’’ and Video 1). Mesothelial rupture initiation was also characterised by the elongation of E-cadherin puncta, and we did not observe any dramatic reduction of E-cadherin at intact regions of the mesothelial cell-cell contacts during rupture initiation (Fig. 1d’’’, Video 1). These E-cadherin fibrillar bridges are not specific to mesothelium cultured on glass, as we observe fibrillar bridges at sites of spheroid invasion when mesothelium is grown on Matrigel (Video 2). Notably, fibrillar bridges and microvoids in polymer crazing are ∼10-20 nanometres, much smaller than the E-cadherin fibrillar bridges and microvoids between cells we observed (Fig. 1d’’ and Video 1). Thus, instead, we termed the formation of E-cadherin fibrillar bridges and microvoids during the initiation of cell-cell junction rupture as cell-cell contact crazing.

We weakened cell-cell contacts to test if these E-cadherin fibrillar bridges help resist rupture. (Fig. 1d, Video 1 & 2). RhoA-Myosin-II mediated contractility supports E-cadherin cell-cell adhesion strength^19-22^. We reduced E-cadherin adhesiveness of the mesothelium by expressing a dominant negative RhoA mutant T19N fused to E-cadherin (Fig. 1e and Supplementary Fig. 1c-f). Strikingly, the E-cadherin fibrillar bridges formed during spheroid invasion were reduced in the mesothelium expressing E-cad-RhoA-T19N mutant (Fig. 1f,g). Although the rate of mesothelial rupture remains unchanged in mesothelium expressing E-cad-RhoA-T19N mutant (Supplementary Fig. 1g), more extensive mesothelial clearance by spheroid invasion was observed (Fig. 1h). Thus, these mesothelial E-cadherin linked fibrillar bridges may resist invasion. Taken together, we observe features of material-based fracture during cell invasion of the mesothelium.

### Tensile fracture of the mesothelium by spheroid engagement

Next, we identify the underlying patterns of motion and mode of mesothelium rupture involved in invasion. To analyze the mesothelium fracturing process, we tagged its apical surface with fluorescent microbeads^23^ and imaged it undergoing spheroid invasion. 500nm beads were coated with wheat-germ-agglutinin for cell surface binding^23^.

A material’s fracture response can be classified as brittle or more ductile according to the extent of its deformation during fracture^24^. A ductile fracture is seen in readily deformable materials and is characterized by extensive plastic deformation before the material finally breaks apart^25^. By contrast, a brittle fracture exhibits little or no plastic deformation before its fracture^25^. Velocity magnitude quantification from particle image velocimetry (PIV) of the WGA-coated beads on the mesothelium apical surface showed significant deformation of the mesothelium (Fig. 1i, Supplementary Fig. 1h). This suggests that the tearing of the mesothelium is akin to ductile rupture.

Generally, there are three principal modes by which fracture/rupture occurs, namely tensile fracture^26,27^ (mode I), shear fracture (mode II) and out-of-plane tearing (mode III) (Fig. 1j). Since cell-cell contact crazes typically form at the plane normal to tensile stress, spheroid invasion should likely induce mesothelial tensile fracture.

Indeed, PIV of the beads on the mesothelium apical surface after spheroid invasion showed a constriction zone next to the region where the crack was initiated (Fig. 1i, Supplementary Fig. 1h). After being pulled from both sides, the small crack opened a hole in the middle of the mesothelium, an indication of a tensile fracture.

Given that cells did not slide past each other upon fracture, it is unlikely that in-plane shear is an important driver of fracture initiation (Fig. 1d). To quantify the deformation nature of the mesothelium, we examined both the normal strain rate and shear strain rate of the mesothelium undergoing fracture (Fig. 1i,k, Supplementary Fig. 1h). Here, the normal strain rate represents the strength of the velocity gradient in the direction of the local velocity, and the shear strain rate represents the strength of the velocity gradient perpendicular to the local velocity^28^. We did not observe a high shear rate near the initial fracturing site (Fig. 1i,k, Supplementary Fig. 1h). Instead, we observed a low shear rate with high expansion velocity, thus corresponding to tensile fracture (Fig. 1i,k, Supplementary Fig. 1h, Methods).

Finally, using lattice light sheet microscopy to achieve a clear cross-section view, we did not observe any out-of-plane tearing (mode III) of the mesothelium caused by the spheroid (Fig. 1l). Protrusions penetrating the mesothelium E-cadherin junctions preceding mesothelial fracture were only observed in ∼6% of the invading spheroids (Supplementary Fig. 1i). While in most of the spheroids, their protrusions were formed after mesothelial fracture. Indeed, lattice light sheet imaging showed that mesothelial cells were forced apart from the apical side first, and such separation then propagated to the basal side, leading to a complete rupture of the mesothelium. Thus, spheroid invasion induces tensile rupture, and we have ruled out other modes of tissue rupture.

### Spheroid leader cell attachment initiates apical constriction and tensile fracture of the mesothelium

We then investigated the cellular mechanism underlying tension-induced mesothelial fracture by spheroids. A large displacement at one side of the microvoid (Fig. 1i) may reflect an apical constriction starting from a mesothelial cell beneath the spheroid as the mesothelium begins to fracture (Fig. 2a). By contrast, such aggressive apical constriction is not detected in other areas of the intact mesothelium away from the fracture site as shown by a more constant cell apical area (Fig. 2b). It is conceivable that apical constriction of a cell could stretch its junctions with neighbouring cells and eventually result in the aforementioned tensile fracture (Fig. 1i). Apical constriction is a cell shape deformation mechanism, promoting tissue remodelling during development^29-32^, that is triggered by increased activation of myosin-II at the apical cortex^21,31,33,34^. Structured illumination super-resolution imaging of myosin-regulatory-light-chain (MRLC)^35^ showed a myosin-II network extending across the apical region of the mesothelial cells (Fig. 2c, c’). In quantitative live-imaging, MRLC intensifies at the junctional and apical cortex region of a mesothelial cell beneath the interacting spheroid (Fig. 2a,d and Video 3). Such intensification of MRLC precedes junction constriction and fracture (Fig. 2d). These observations support the notion that constriction of apical mesothelial myosin-II cortex pulls on E-cadherin junctions, eventually leading to initiation of mesothelial fracture.

**Figure 2.**
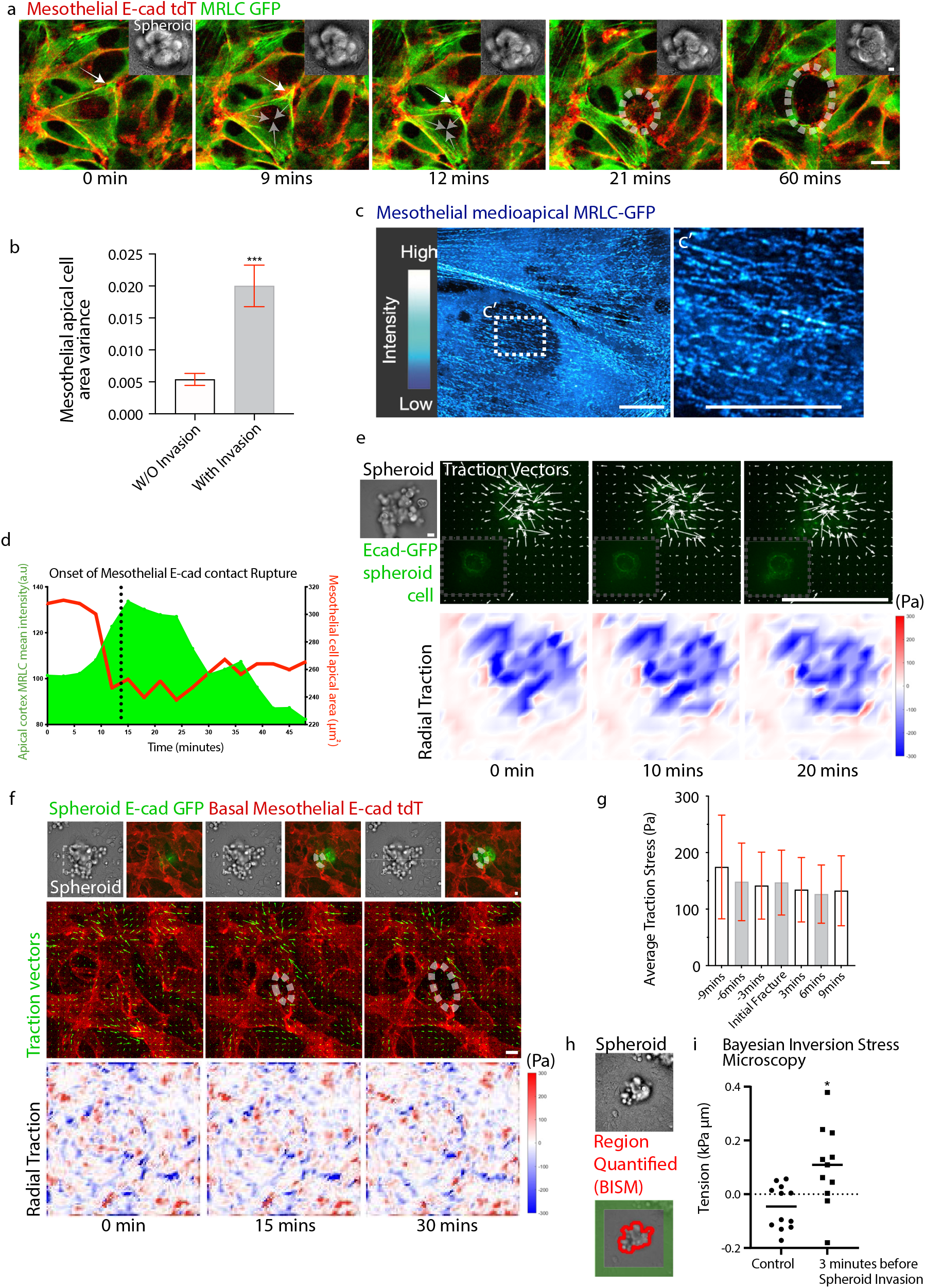
Initiation of fracture from apical constriction of the mesothelial cell. (a,b,c,d) Apical constriction precedes fracture initiation of the mesothelium. (a) Time-lapse images of the spheroid (brightfield, top right) invading mesothelium expressing E-cad tdT and MRLC-GFP. Intensification of MRLC filaments is observed as apical constriction of the mesothelial cell led to subsequent fracture initiation of the mesothelium. (b) Variance quantification of mesothelial apical cell area beneath spheroid during fracture compared to areas not undergoing fracture. (c) Network of myosin-II filaments spans the apical region of the mesothelium. Structured illumination microscopy of mesothelium expressing MRLC-GFP, with a detailed view in (c’) (d) Myosin-II filaments intensification precedes apical constriction. Representative temporal changes in myosin-II filaments intensity at the apical cortex and size of mesothelial cell apical area. (e) Spheroid leader cell contracts basal substrate inwards. Time-lapse images of radial traction stress of E-cad GFP expressing spheroid leader cell attaching onto the PDMS substrate. Heat maps in (e,f) shows the direction and magnitude of radial traction stress. The colour scale bar indicates the magnitude of the outward (red) and inward (blue) radial traction stress from the mesothelial fracture site. (f,g) Traction force of mesothelial cells did not exhibit obvious changes upon fracture. Time-lapse images (f) and quantification of fracture site average cellular traction stress (g) of E-cad tdT expressing mesothelium when invaded by spheroid expressing E-cad GFP. The discontinuous line in (f) indicates the area of the fracture. (h,i) An increase in mesothelial tensile stresses at regions surrounding the invading spheroid prior to invasion. Comparison of isotropic tensile stress at a 10-micron boundary (h, red) surrounding invading spheroids prior to invasion and control which are spheroids that fail to invade (i). For green regions highlighted in (h), refer to Supplementary Fig. 2b. All data are means ± SEM from n=7 cells from at least 3 independent experiments, except for (b), which is from 4 independent experiments, *p < 0.05, analysed with Mann-Whitney U test. Scale bars, 5µm.

Besides the apical constriction mechanism, we use traction force microscopy (TFM) to analyse the contribution of in-plane traction-mediate-migration of mesothelial cells during fracture^23,26^. First, we find that the spheroid leader cell tends to contract the Polydimethylsiloxane (PDMS) substrate in the absence of mesothelium (Fig. 2e). Such inward contraction from spheroid cells is well poised to activate the inward contraction/apical constriction on contacting mesothelial cells.

Next, we introduced a spheroid on the mesothelium grown on PDMS but failed to detect an enhancement in the mesothelial cell traction force on the PDMS substrate (Fig. 2f,g) or any change in radial traction stress of mesothelial cells upon fracture initiation (Fig. 2f, Supplementary Fig. 2a). Unexpectedly, in contrast to previous qualitative observations, we did not observe traction-mediated lateral migration of mesothelial cells away from the invading spheroid during fracture initiation^11^. Notably, our finding reinforces the conclusion that mesothelial fracture ductility has absorbed and dissipated forces exerted by the spheroid (Fig. 1j,k,l), preventing any obvious changes in the patterns of mesothelium traction forces. Although obvious changes in the patterns of mesothelium traction forces are undetected, Bayesian inversion stress microscopy analysis^36^ identifies an increase in supracellular mesothelial tensile stresses surrounding the spheroid before invasion (Fig. 2h,i, Supplementary Fig. 2b). The increase in mesothelial tensile stresses is likely due to the reorganization of apical actomyosin filaments (Supplementary Fig. 2c-g). Taken together, this reinforces that tensile stresses, likely due to apical constriction of the mesothelium, are responsible for invasion-induced fracture.

### Intercellular and apical integrins trigger mesothelial fracture and collective cell invasion

We then look deeper into the molecular mechanisms underlying the forces driving invasion. We hypothesize that the spheroid-mesothelium interactions trigger mesothelial apical constriction. It was shown that mesothelial clearance is initiated by ovarian tumour attachment to the mesothelium via integrin alpha-5^11,12^, but the organization of intercellular integrin-matrix complexes in a 3D microenvironment remains unclear. We imaged Paxillin - a key component of the integrin matrix adhesion at the spheroid-mesothelium contact region to visualise the intercellular integrin-matrix complexes. Strikingly, when they attached to the mesothelium, spheroids expressing Paxillin-GFP exhibited strands of Paxillin at the spheroid-mesothelium interface (Fig. 3a, Supplementary Fig. 2h), which grew from smaller initial spots (Fig. 3b). The mesothelium tends to rupture after a spheroid’s leader cell spreads on a mesothelial cell (Supplementary Fig. 2i). These strands of Paxillin morphologically resemble the integrin adhesion complexes seen when cells spread on glass^37^. We observed that mesothelial rupture occurred after spreading of a spheroid’s leader cell on the mesothelium in ∼70% of invasion cases (Supplementary Fig. 2j).

**Figure 3.**
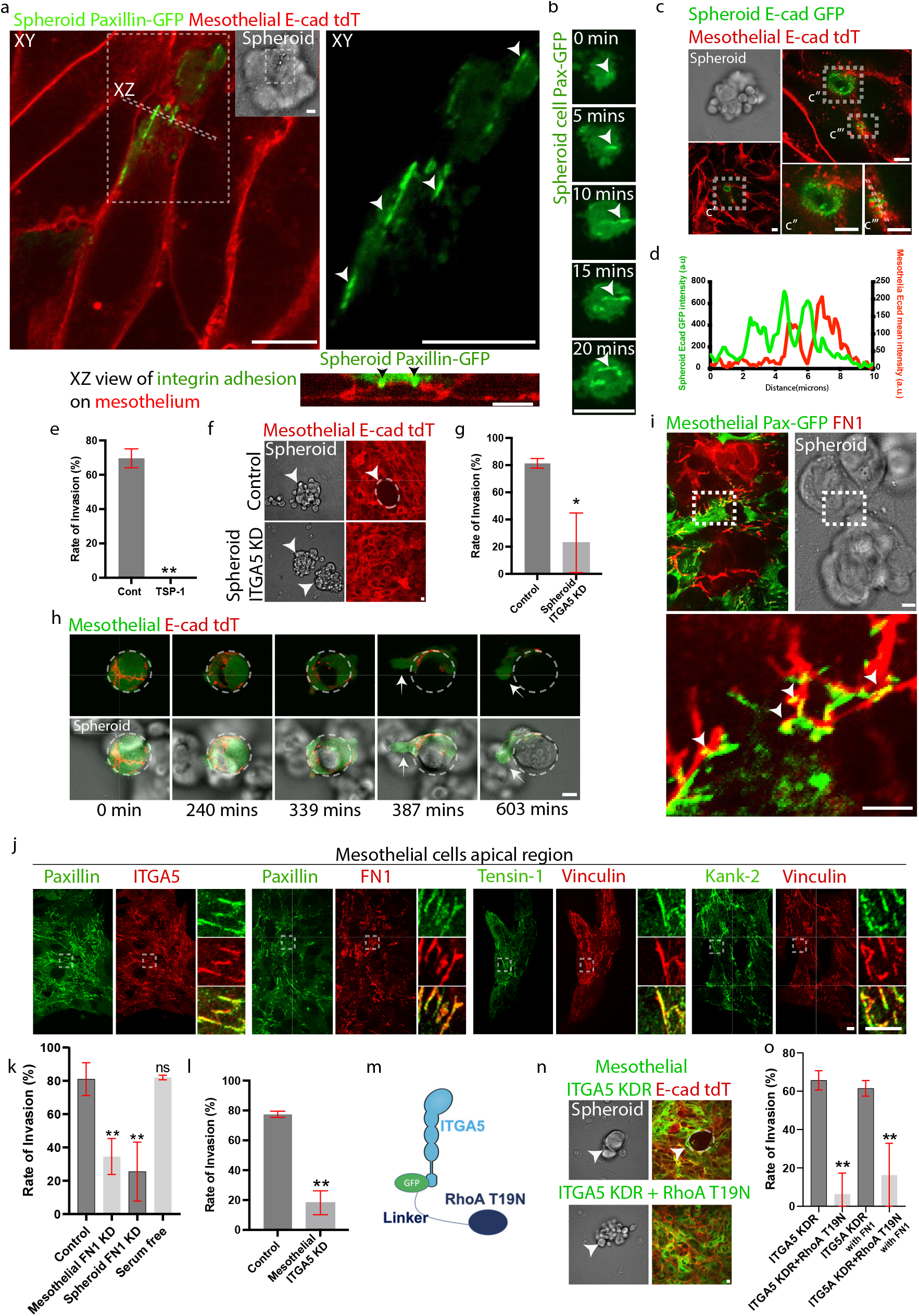
Intercellular integrin adhesion and fibronectin matrix complexes connect the spheroid to the mesothelium. (a) Strands of integrin adhesion clusters at the spheroid-mesothelium interface from spheroid expressing Paxillin GFP with indicated XZ representative views (bottom). (b) The spheroid leader cell’s Paxillin GFP clusters grow gradually at the spheroid-mesothelium interface. The cell that is first attached to the mesothelium is defined as the invasion leader cell. Time-lapse imaging of Paxillin GFP at the spheroid-mesothelium interface. (c) Cadherin at the interface between spheroid (top left, brightfield) expressing E-cad GFP and mesothelium expressing E-cad tdTomato does not co-localize, with detailed views of c’ in c’’ and c’’’. (d) Representative fluorescence line-scan analysis of spheroid E-cad GFP and mesothelium E-cad tdTomato in c’’’ shows a lack of homophilic E-cadherin interactions. (e) Thrombospondin-1 peptide treatment, blocking integrin-based fibronectin attachment, prevents spheroid-induced mesothelial fracture. Rate of spheroid-induced fracture in control compared to thrombospondin-1 peptide treated cultures. (f,g) Spheroid integrin knockdown blocks fracture. Representative images (f) and quantification of the spheroid (g) invading the mesothelium in control compared to ITGA5 knockdown in the spheroid. (h) Mesothelial cells associating with spheroids at non-adhesive regions of micropatterned glass. Time-lapse images of spheroid displacing mesothelial doublets from round adhesive fibronectin patterns into non-adhesive zones. (i) Co-localization of fibronectin with apical mesothelial integrin clusters. Immunofluorescence images of immunostained fibronectin and mesothelium expressing Paxillin-GFP with detailed views (bottom). (j) Mesothelial apical integrin adhesion complex without spheroid engagement. Immunofluorescence images of mesothelium apical region immunostained for Paxillin, ITGA5, FN1, Kank-2 and Vinculin. (k,l) FN1 and ITGA5 from the mesothelium and spheroid are required for spheroid-induced mesothelial fracture. Rate of spheroid-induced fracture in control compared to either mesothelium FN1 knockdown, spheroid FN1 knockdown, serum-free media condition (k) or ITGA5 knockdown (l). (m) Design of ITGA5-GFP fused to RhoA dominant negative T19N mutant for expression in the mesothelium. (n,o) Spheroid-induced fracture requires activation of contractility at sites of mesothelial ITGA5 adhesion. Representative images (n) and quantification (o) of spheroid-induced mesothelial fracture rate in control mesothelium expressing ITGA5 knockdown with reconstitution of ITGA5-GFP compared to the reconstitution of ITGA5-GFP fused to RhoA dominant negative T19N mutant in the mesothelium. All data are means ± SEM from at least with ∼50 spheroid from n=3 independent experiments, *p < 0.05, **p < 0.01; analysed with unpaired t-test. Scale bars, 5µm.

As E-cadherin interactions between the epithelial assemblies^8,19,22,33,38-43^ were undetected with co-localization analysis between E-cad GFP spheroid and E-cad tdT mesothelium (Fig. 3c,d), we continued to analyse the role of intercellular integrin instead. Knocking down α5 integrin and using a thrombospondin-1 (TSP-1) peptide^44^ to specifically block fibronectin-based integrin adhesion of cells successfully blocked mesothelial fracture by the spheroid (Fig. 3e-g, Supplementary Fig. 2j,k), validating the role of α5 integrin in the spheroid-mesothelium attachment.

Adhesion receptors such as integrins could be present at the apical surface of mesothelial cells, mediating their attachment to the spheroid. This is supported by our finding that mesothelial cells detached from the adhesive (i.e. fibronectin-coated) round micropattern can be associated with the spheroid at non-adhesive region of the substrate (Fig. 3h). Whereas in controls, mesothelial cells remained attached on the adhesive round micropattern (Supplementary Fig. 2l). Additionally, this observation was consistent with our TFM data, together suggesting that mesothelial cell’s outward migration away from the spheroid by basal traction is dispensable for mesothelial fracture.

Strikingly, strands of Paxillin were localized at the apical side of mesothelial cells and exhibited co-localization with immunostained fibronectin (Fig. 3i). Even in the absence of spheroid attachment to the apical side of the mesothelium, α5 integrin and other binding partners of the integrin complex, including Paxillin, kank2, tensin1, fibronectin and vinculin, co-localized at sites of apically localized integrin-based adhesions (Fig. 3j). Notably, the mesothelial apical integrin that we have identified here, although not visualised previously, is reported to functionally mediate apical constriction in the embryonic Drosophila epithelium^45,46^.

It was suggested that integrin and Talin-dependent activation of non-muscle myosin-II was required for an uncoupling mechanism to separate the mesothelial cells from fibronectin fibrils to drive mesothelial cell migration away from the spheroid during invasion^11,12^. However, instead of such an uncoupling mechanism, the mesothelial apical fibronectin network undergoes fracture as the spheroid invades (Video 4). Thus, our data shows that ECM and mesothelium barrier fracturing mechanism is the dominant invasion mechanism.

Furthermore, mesothelial α5 integrin or mesothelial/spheroid fibronectin knockdown was sufficient to block spheroid-induced fracture (Fig. 3k,l, Supplementary Fig. 2m-p and Video 5). Whereas fibronectin from the media is not required for invasion, as invasion remains robust in serum-free media experimental condition (Fig. 3k, Supplementary Fig. 2n). Of note, knockdown of α5 integrin primarily depleted apical integrins with a subtle reduction of basal integrins (Supplementary Fig. 2o). Thus, the mesothelium’s apical integrin adhesion complex and spheroid fibronectin are required for spheroid invasion.

Our evidence suggests that mesothelial apical constriction from intercellular integrin engagement initiates the mesothelial tensile fracture. To directly test this notion, we blocked spheroid-induced mesothelial apical constriction by linking a dominant-negative RhoA mutant to α5 integrin (Fig. 3m and Supplementary Fig. 2q-t). This significantly reduced spheroid-induced mesothelial fractures (Fig. 3n,o, Supplementary Fig. 2u,v, and Video 5). Although RhoA inhibition reduces mesothelial apical fibronectin fibril assembly (Supplementary Fig. 2w,x), the restoration of fibronectin fibrils by introducing exogenous fibronectin-1 fails to rescue the reduction of invasion caused by RhoA inhibition (Fig. 3n,o, Video 6). Thus, intercellular integrin engagement induces mesothelial fracture, in part by apical myosin-II activation, is the key molecular mechanism driving invasion.

### Spheroid invasion of the mesothelium did not involve induction of jamming transition

We then addressed how spheroid multicellular dynamics affect invasion. Invading spheroids break through the mesothelium and then attach to the submesothelial substrate to prevent anoikis of spheroid cells^47^(Fig. 4a). Thus, we assess the critical relationship between spheroid cell dynamics and mesothelial fracture. We quantitatively compared the cellular movement pattern in (1) invading spheroids with (2) non-invading spheroids due to ITGA5 knockdown in the mesothelium. We obtained tracking data of the spheroid’s cell nuclei by deep-learning segmentation^48,49^ and IMARIS tracking (Fig. 4b,c, Supplementary Fig. 3a). In invading spheroids, there was a spreading of cells at the bottom layer, as well as an overall bias in cell movement towards the mesothelium base even before mesothelial fracture (Fig. 4c,d, Video 7). By contrast, the cell movement in non-invading spheroids has no particular direction (Fig. 4d,e). We then assess whether directed cell motion is involved in spheroid invasion by quantifying the persistency of cell velocity (Methods). Our results suggest that cells in non-invading spheroids move less persistent than those in invading spheroids (Fig. 4f).

**Figure 4.**
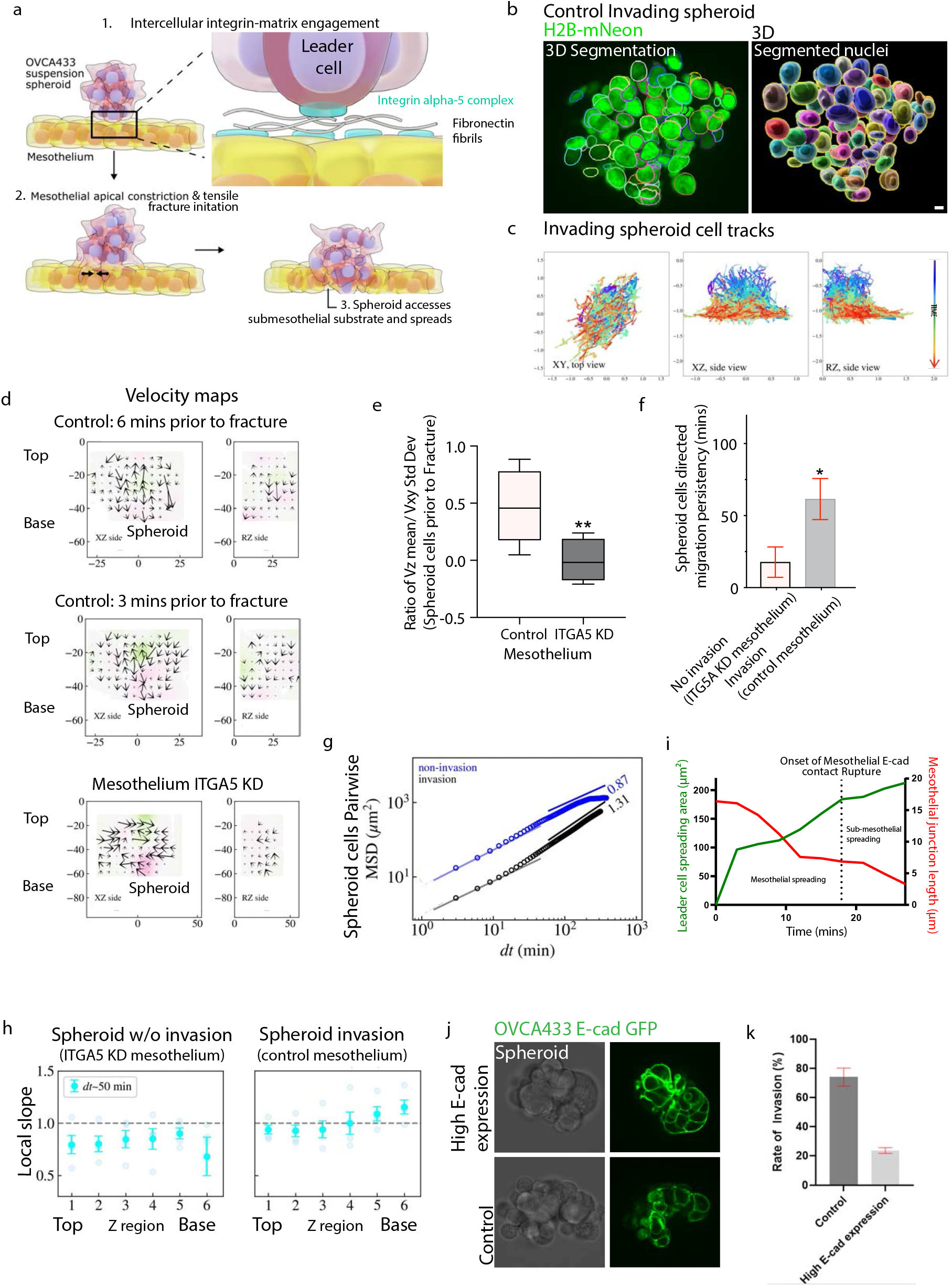
Jamming transition did not induce the spheroid invasion of the mesothelium. (a) Model for spheroid-induced mesothelial tensile fracture (b) Representative images of an OVCA433 spheroid labeled with H2B-mNeon segmented by 3D StarDist (left) and a 3D rendering of the segmented nuclei. (c) Representative 2D projection of a 3D cell trajectories in spheroid invading the mesothelium over time: (left) in XY axis, projected from the top, (middle) map in XZ axis projected from the side and (right) map in Radial Z (RZ) axis, a radial projection of spheroid assuming the rotational symmetry around the axis at the in-plane center of the spheroid. (d) Spheroid cells exhibit motility bias towards the mesothelium prior to mesothelial fracture. Representative XZ and RZ side view velocity maps of a spheroid 6 and 3 minutes before mesothelial fracture compared to spheroids that will not induce fracture due to ITGA5 knockdown of the mesothelium. (e) Mesothelium ITGA5 is required for spheroid cells motility bias towards the mesothelium. The ratio of spheroid cells Z velocity mean over standard deviation of XY velocities distribution of spheroid cells on control mesothelium compared to spheroid cells on ITGA5 knockdown mesothelium. (f) Persistency of directed spheroid cell migration (see Methods) is higher in spheroid invading the mesothelium compared to spheroid without invasion due to ITGA5 knockdown in the mesothelium. Quantification of spheroid cells velocity auto-correlation persistency in spheroid with or without invasion. (g, h) Pair-wise mean square displacement (MSD) (g) (see Methods) of representative samples for spheroid with or without invasion. (h) Local slope of MSD value obtained from the pair-wise mean square displacement graph for six selected regions from top to base of spheroids (see Methods) for two different conditions. (i) Shrinkage of mesothelial cell-cell junctions and spreading of leader cell on mesothelium precedes fracture. Representative temporal changes in leader cell spreading area at mesothelial and sub-mesothelial regions is in green, and mesothelial junction length is in red. Dotted line indicates the onset of mesothelial cell-cell contact rupture. (j,k) Spheroids expressing higher levels of E-cadherin have reduced rate of invasion. Representative E-cad GFP expressing spheroid images (j) and quantification of the spheroid (k) invading the mesothelium in control compared to higher E-cadherin levels expressing spheroid. All data are means ± SEM from at least n=4 independent experiments, *p < 0.05, **p < 0.01, ****p < 0:0001; analysed with Mann-Whitney U test. Scale bars, 5µm.

Jamming transition is a common trigger of collective cell migration, involving the transition of multicellular assemblies from a jammed to fluid state^50-53^. To quantify the involvement of jamming transition, we then quantify the mean-square displacement (MSD) of cell nucleus centroids obtained from AI segmentation and cell tracking. To exclude the effect of a global drift of the spheroids, we adopted pair-wise MSD for our system (see Methods). We found that at a small timescale of about 10 mins, both non-invading and invading spheroids show roughly diffusive movement with MSD linearly growing with lag time, while on the timescale larger than 50 minutes, MSD grows with lag time super-linearly for invading and free-spreading spheroids and sub-linearly in non-invading spheroid (Fig. 4g,h). The short-time diffusive movement of cells suggests that the spheroid is in a fluid state^36,37,38^ even when the spheroid cannot invade due to the lack of ITGA5 expression in the mesothelium. Our 3D cell tracking and MSD analysis indicate that cancer cells are in an unjammed state even when confined as a spheroid with intact E-cadherin adhesion. Thus, the triggering of jamming transition is not a key driver for cancer spheroid invasion and spreading.

Instead, the whole spheroid could be considered as a multicellular droplet held by an effective tensile surface. The long-time sub-linearity of MSD with non-invading spheroids indicates the caging effect brought by the boundary of this effective surface, whereas, in invading spheroids, the super-linearity of MSD on the larger timescale implies the existence of some external force that guides cells in persistent and directed movement. Thus, this external force likely originates from wetting at the intercellular integrin cell-cell contact between the spheroid leader cell and the mesothelium. Indeed, α5 integrin and fibronectin are required for the invasion. Moreover, the spreading of the spheroid leader cell coincides with the shrinkage of the underlying mesothelial cell-cell junction preceding mesothelium fracture (Fig. 4i). Unexpectedly, spheroid invasion is independent of jamming transition induction.

### Theoretical modelling of migration and tissue invasion

To elucidate the underlying physical mechanisms behind the observed experimental phenomena, we developed a theoretical framework grounded in the concept of active wetting (Methods). Specifically, we modelled the spreading and invasion of leader spheroid cells as a wetting-driven interfacial displacement process, in which the system’s free energy is determined by interfacial energy and polarization work (Eq. 1). The interfacial energy, in turn, evolves dynamically according to interfacial adhesion and cortical tension, as governed by Eqs. (2∼4).

Our theoretical model predicts the critical conditions under which spheroid cell invasion occurs. A physical process can only proceed spontaneously when the change in free energy is negative (Fig. 5a), thereby providing a clear boundary between invasive and non-invasive regimes (Fig. 5b,c). The theoretical analysis indicates that the critical condition for spontaneous invasion is characterized by a 1/2 power law (Eq. 8) between the spheroid-mesothelial integrin adhesion strength (*W*_*I*0_, Methods) and the activated contractility of mesothelial cells (described by the coupling strength β, Methods). Moreover, due to the apical constriction of mesothelial cells, the effective adhesion length between the two cell types is substantially reduced, which in turn increases the required integrin-mediated adhesion strength for effective invasion. As expected, the critical condition also shows that polarization of the spheroid cells toward the mesothelial layer lowers the energy barrier and facilitates invasion, while an increase in the thickness of the barrier cell layer impedes it (Fig. 5b,c).

**Figure 5.**
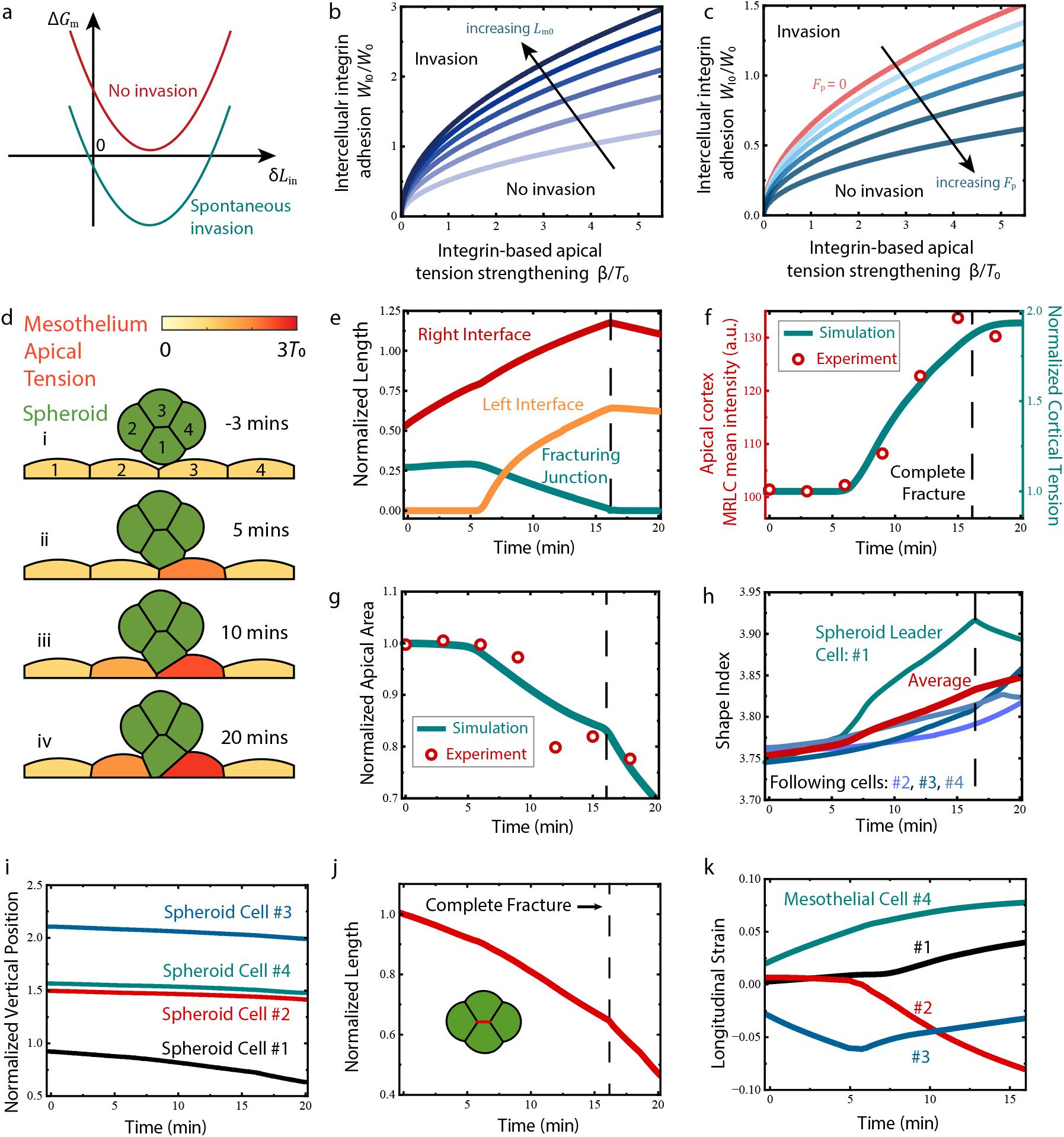
Theoretical predictions and numerical simulations of wetting-induced invasion. (a-c) Theoretical predictions for spontaneous fracture initiation and spheroid invasion. (a) Schematics for the relationship between active wetting (increasing *δL*_in_) and free energy change, where a spontaneous invasion sets the condition as Δ*G*_m_ ≤ 0. (b) Increasing mesothelial monolayer thickness (represented by increasing *L*_m0_) significantly pushes the critical invasion conditions toward higher intercellular integrin adhesion. (c) Spheroid polarization promotes cell invasion and lowers the requirements of *W*_I0_ for complete fracture initiation. (d-k) Simulated invasion process agrees with experimental results when the intercellular adhesion is comparable to the mesothelial E-cadherin adhesion, i.e. when W_I0_ ⁄W_0_ =0.9, W_0_⁄ *T*_0_ =1.2, *β⁄ T*_0_ =1.5. (d) Intercellular integrin adhesion drives spheroid invasion and mesothelial fracture. Stages of collective invasion include [i] initial state, [ii] establishment of contact between the leader spheroid cell and the 2^nd^ mesothelium cell, [iii] invasion, and [iv] complete mesothelial fracture. To make a direct comparison with experimental results, we set the initial time as −3 mins. (e) Complete fracture occurs ∼10 mins after the spheroid leader cell comes into contact with the 2^nd^ mesothelial cell. The right interface refers to the interface between the leader spheroid cell #1 and the #3 mesothelial cell, the left interface refers to the interface between the leader spheroid cell #1 and the #2 mesothelial cell, and the fracturing junction refers to the junction between #2 and #3 mesothelial cells labelled in (d). (f) Simulated apical tension of the #2 mesothelial cell is in qualitative agreement with the observed MRLC results. The computed tension is normalized by the static mesothelial tension *T*_0_. (g) Simulated apical area of the #2 mesothelial cell fits well with the experimentally measured apical cell area. The apical area is calculated by the square of the simulated apical cell length and is normalized by the initial area. (h) Spheroid cells show T1 transition during the invasion process. The shape index (individual and average) of the spheroid cells is used to quantitively evaluate the rearrangement of the spheroid. (i) Simulated collective spheroid cell motion towards the direction of the mesothelium (negative Z direction). (j) Central spheroid junction shrinkage indicates striking deformation and intercalation of spheroid cells. (k) Longitudinal strain of the mesothelial cells. Longitudinal strain is computed by the dynamic change of cell length.

To directly compare theoretical predictions with experimental results, we conducted numerical simulations of the invasion process using the proposed theoretical framework in conjunction with the active foam model^54^ (Supplementary Fig. 3b, Methods). Key simulation parameters were informed by experimental observations and cortical tension measurements by atomic force microscopy (AFM) (Supplementary Fig. 3c,d, Methods).

The simulations (Fig. 5d-k) reproduced all key experimental observations, including the integrin-mediated extension of the spheroid-mesothelial interface, enhanced cortical tension due to active mesothelial contraction, E-cadherin junction fracture between mesothelial cells in contact with spheroid cells, and internal cellular rearrangements within the invading spheroid cell (Fig. 5d,e, Video 8, top left). The accuracy of our model was further validated by comparing simulated and experimentally observed apical constriction areas and apical tension in mesothelial cells (Fig. 5f,g). In particular, by tracking the changes in spheroid cell shape (quantified by the shape index *I*_*S*_, a key descriptor of cell morphology and intercellular junction lengths (Fig. 5h), our simulation identified that spheroid cells undergo prominent T1 transition during initiation of mesothelial invasion. This leads to super-diffusive motion of the spheroid mass (Fig. 5i,j), consistent with experimental observations (Fig.4d,e,g).

Numerical modelling enabled us to quantitatively control the parameters of invasion; thus, we seek to identify key factors affecting the invasion process. As a direct driver of invasion, weakened integrin adhesion between cells slowed down spheroid rearrangement and ultimately abolished the spheroid’s ability to invade the barrier layer (Fig. 6a-e; Video 8, top right). Conversely, increasing integrin adhesion promoted rapid internal rearrangement of spheroid cells and accelerated rupture of the mesothelium (Fig. 6f-j; Video 8, bottom left). We did not observe an obvious effect of reduced E-cadherin strength in mesothelial cells on collective invasion (Supplementary Fig. 3e-h, Video 8, bottom right). Nonetheless, this reduction caused greater deformation of neighboring mesothelial cells during invasion (Fig. 5k, Supplementary Fig. 3i), which may explain the experimentally observed increased mesothelial clearance upon E-cadherin adhesion weakening (Fig. 1h).

**Figure 6.**
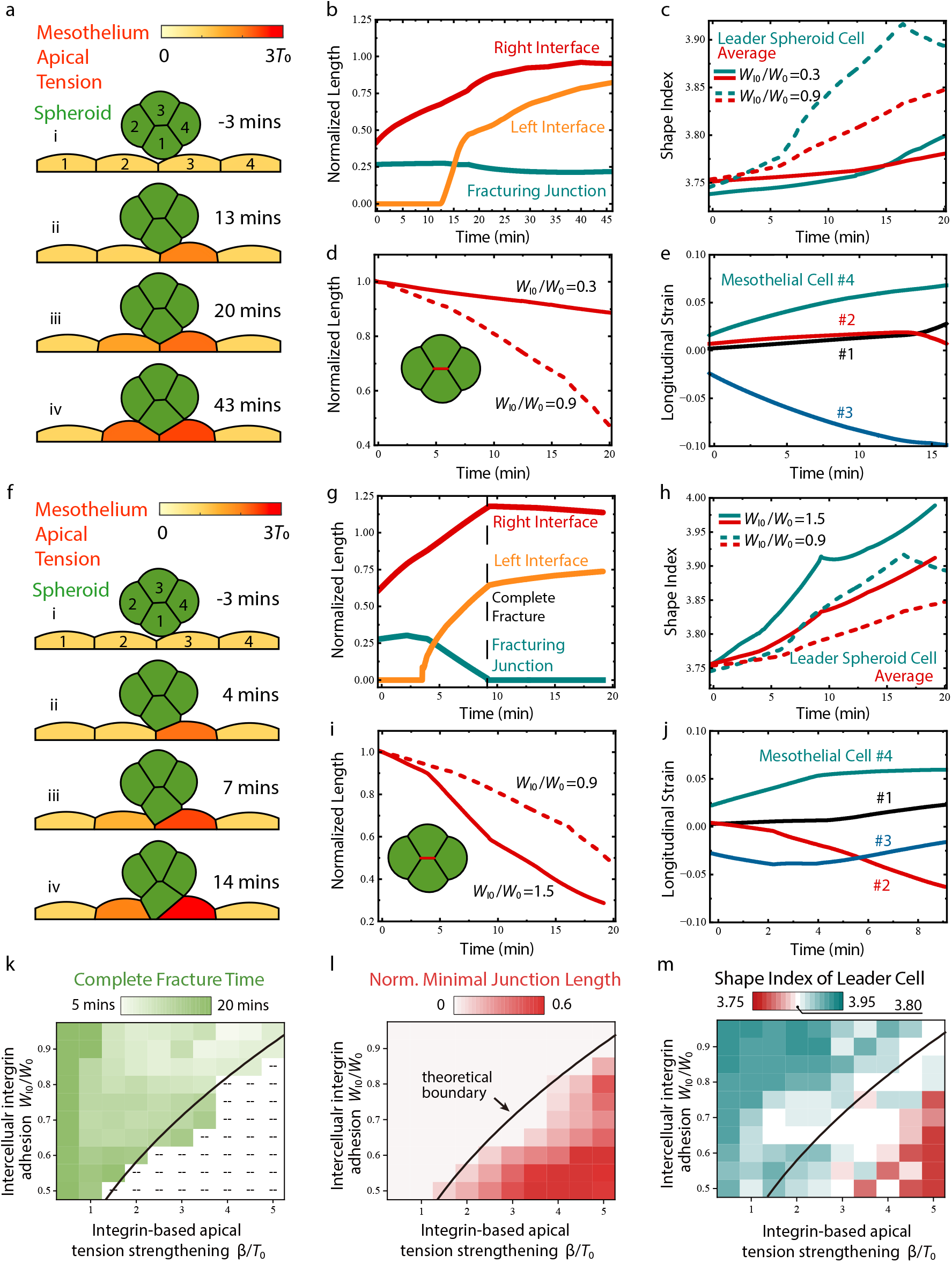
Extensive simulations show apical constriction level and adhesion strength as key factors affecting invasion dynamics. (a-e) Low integrin adhesion strength (*W*_I0_⁄*W*_0_ =0.3, *W*_0_⁄ *T*_0_ =1.2, *β*⁄*T*_0_ =1.5) prevents spheroid T1 transition and leads to incomplete mesothelial fracture. (a) Process of incomplete fracture includes [i] initial state, [ii] contacting between the leader spheroid cell and the 2^nd^ mesothelium cell, [iii] further invasion, and [iv] complete mesothelial fracture nucleation. The initial time is set as −3 mins. (b) Fracturing junction reaches its minimal length ∼25 minutes after contacting the 2^nd^ mesothelial cell, indicating an incomplete fracture at low integrin adhesion strength. The right interface refers to the interface between the leader spheroid cell (#1) and the #3 mesothelial cell, the left interface refers to the interface between the leader spheroid cell (#1) and the #2 mesothelial cell, and the fracturing junction refers to the junction between #2 and #3 mesothelial cells. (c) Low integrin adhesion strength prevents T1 transition of spheroid cells during invasion. (d) Low integrin adhesion strength results in slow central junction shrinkage and thus prevents T1 transition in the spheroid. (e) Low integrin adhesion strength changes the distribution of mesothelial strain in adjacent cells. (f-j) High integrin adhesion strength (W_I0_⁄W_0_ =1.5, W_0_⁄ *T*_0_ =1.2, *β* ⁄ *T*_0_ =1.5) results in striking spheroid T1 transition and mesothelial fracture. (f) Dynamic process of rapid fracture includes [i] initial state, [ii] contacting between the leader spheroid cell and the 2^nd^ mesothelium cell, [iii] further invasion, and [iv] complete mesothelial fracture nucleation. The initial time is set as −3 mins. (g) Complete fracture occurs ∼6 mins after the leader cancer cell comes into contact with the 2^nd^ mesothelial cell. The right interface refers to the interface between the leader spheroid cell #1 and the #3 mesothelial cell, the left interface refers to the interface between the leader spheroid cell #1 and the #2 mesothelial cell, and the fracturing junction refers to the junction between #2 and #3 mesothelial cells. (h) High intercellular integrin adhesion strength leads to striking T1 transition of spheroid cells during invasion. (i) High integrin adhesion strength results in quick central junction shrinkage and, thus, striking T1 transition in the spheroid. (j) Slightly smaller mesothelial deformation was observed by increasing intercellular integrin adhesion strength. (k-m) Mesothelial fracture and spheroid rearrangement are strongly interrelated during invasion. The simulated critical boundary for the occurrence of mesothelial fracture agrees well with the prediction (black curves) given by Eq. (5). Complete mesothelial fracture and spheroid rearrangement only occur at low to medium apical contractility and high integrin adhesion. Low integrin adhesion strength or high apical contractility leads to lower spheroid T1 transition and incomplete fracture. (k) Complete fracture time as a function of the integrin adhesion strength and apical contractility. Rapid invasion occurs at medium apical contractility and strong integrin adhesion strength. No data is at the bottom right corner since no complete mesothelial fracture occurs in this region. The time taken for fracture is calculated as the period between the contact of the leader spheroid cell with the 2^nd^ mesothelial cell and the complete rupture of the mesothelial junction. (l) High apical contractility with low intercellular integrin adhesion strength leads to incomplete fracture, i.e., a non-vanishing minimal junction length (in the lower right region) between #2 and #3 mesothelial cells labeled in (Fig. 5d). The minimal fracturing junction length is normalized by its initial length. (m) Spheroid cell rearrangement is regulated by intercellular integrin adhesion and apical contractility. The shape index reflects the deformation of the leader spheroid cell with respect to different intercellular integrin adhesion strength and apical contractility. Here, the shape indices are taken when the fracturing junction length reaches 0 (for cases where mesothelium is completely fractured) or when the fracturing junction length reaches its minimum for cases where there is no complete mesothelial fracture.

Finally, we performed simulations to evaluate the impact of mesothelial cell contractility and summarized the results alongside those involving intercellular adhesion strength in phase diagrams (Fig. 6k,l,m, Supplementary Fig. 3j). Specifically, moderate enhancement of apical constriction in barrier cells accelerated invasion, whereas overtly high constriction effectively suppressed both spheroid invasion and junctional rupture (Fig. 6k). Therefore, invasion occurs only within a defined region of the phase diagram characterized by high integrin adhesion strength and low-to-moderate barrier cell contractility. Moreover, consistent with apical constriction without tissue fracture during embryonic development, our theory and numerical simulations predict that when the intercellular integrin-mediated adhesion strength is weak (or absent), apical constriction alone does not drive the rupture of the barrier cell monolayer (Fig. 5b,c, Fig. 6k,l). The invasion–non-invasion boundary predicted by simulations aligned closely with theoretical predictions (Fig. 6k,l), further confirming that spheroid cell invasion is governed by an interfacial displacement process driven by active tissue wetting. Similarly, due to the competition between integrin adhesion–mediated wetting and mesothelial apical constriction–driven dewetting, the spheroid cells exhibited morphological transitions between wetting and dewetting states, as evidenced by a significant difference in shape index between the invasion and non-invasion regions in the phase figures (Fig. 6m, Supplementary Fig. 3j).

## Discussion

Prevailing models have attributed collective invasion to unjamming. We challenge this interpretation. Our analyses show that spheroid cells maintaining E-cadherin–mediated contacts are already in an unjammed state, as indicated by mean squared displacement measurements. Although similar dynamic signatures have been reported in other collective invasion studies and interpreted within a jamming framework,^52,55-57^ such interpretations are imprecise because a solid-to-fluid state transition is not evident. In condensed matter physics, unjamming refers specifically to a glass-like solid becoming fluid as packing constraints are relieved. By contrast, the longer-timescale (∼50 mins) dispersal dynamics observed here, likely reflecting multicellular spheroid spreading, indicates a liquid-to-gas–like transition rather than a solid-to-liquid unjamming.

Moreover, spreading of a single leader cell is sufficient to initiate collective invasion and barrier fracture, indicating that invasion does not require a multicellular phase transition such as unjamming. Taken together, these observations demonstrate that unjamming neither precedes nor drives tissue invasion.

What, then, initiates tissue invasion?

A previous model proposed that spheroid protrusion and dissociation of surface fibronectin fibrils from the mesothelium drive outward mesothelial migration to facilitate invasion.^11^ By contrast, higher-resolution quantitative imaging shows that invasion is initiated through contractility-driven fracture of the mesothelial barrier, resembling material “crazing”.^25^ This barrier failure arises from RhoA-dependent forces transmitted through spheroid integrins to mesothelial apical integrins via the intervening fibronectin matrix, triggering mesothelial apical constriction and ultimately tensile rupture of cell–cell junctions. Together, these results indicate that invasion is driven not by protrusive pushing, fibronectin uncoupling, or jamming transitions, but by contractility-induced tensile fracture of the barrier, enabling subsequent spheroid spreading and mesothelial clearance.

To establish a quantitative framework for this process, we applied wetting theory to model spheroid–tissue interactions. While classical wetting describes passive interfacial energy balance, our findings show that barrier cell contractility actively modulates interfacial energy beyond fluctuations arising from junctional dynamics.^58^ Mesothelial apical constriction, coordinated with intercellular integrin mediated adhesion, exerts dual effects on invasion: moderate integrin based contractility promotes invasion, whereas excessive contractility suppresses it through negative feedback that limits further integrin-mediated adhesion growth. Thus, invasion dynamics are governed not solely by properties of the invading spheroid but by active mechanical regulation from the barrier tissue. This active wetting framework quantitatively predicts interfacial invasion behavior and highlights the tissue barrier as a critical mechanical regulator of tissue invasion. Notably, this conclusion is consistent with prior work demonstrating that myoepithelial barrier cells can restrain and recapture invasive breast cancer spheroids^59^.

In conclusion, rather than invasion driven by protrusive pushing or induction of a jamming transition, we demonstrate that coordinated subcellular contractility in both invading cells and the tissue barrier drives invasion through tensile fracture. These findings challenge prevailing paradigms and establish the tissue barrier as an active mechanical regulator of tissue invasion.

### Limitations of the study

First, whether the mechanism we identify is a key driver of metastatic colonization in cancer patients remains to be determined. Second, consistent with the inward contraction observed at the mesothelial surface prior to invasion, we show that spheroid leader cell spreading induces inward contraction of the TFM PDMS substrate. However, the PDMS substrate used for traction force microscopy does not necessarily recapitulate the mechanical behavior of the mesothelial apical surface.

## Resource availability

### Lead contact

Further information on resources reported in this study should be directed to Yuan Lin (theory) and the lead contact, Selwin K. Wu

### Materials availability

Plasmids generated in this study are available on request. Immortalized non-GFP expressing ZT cells created for this study are available on request. Request and information on the resources and equipment used and reported in this study should be directed to Celestine Z. Ho and will be fulfilled by Boon Chuan Low.

### Data and code availability

The scripts generated in this study have been deposited at our GitHub repository: https://github.com/selwinwu/morphogenesis.git and are publicly available as of the date of publication. The specific version of this resource is also archived in Zenodo under DOI 10.5281/zenodo.16737385.

## Data availability

The authors declare that the data supporting this study’s findings are available within the paper and its supplementary files. The raw data generated in this study are available from the corresponding authors upon reasonable request.

## Code availability

The scripts generated in this study are available from https://github.com/selwinwu/morphogenesis.git

## Acknowledgments

We gratefully thank Ernest Poon, Phoebe Lim, Daniella Lawther, Shiwei Beh, Lee Kang Wei, Hui Ting Ong, Chaoyu Fu, Ti Weng Chew, Xingyu Xia, T Thivakar, Soumya Bunk, Thuan Beng Saw, Jennifer Young, Erik Sahai, Robert Parton and Peter Friedl for their input and discussions during this project. Ivan Yow for providing PDMS dishes for TFM and helping with TFM setup and discussions. Raymond Woo and Samuel Ko for Zeiss lattice light sheet microscopy setup and imaging support. Wei Jia Goh for discussions and suggestions to use 3D StarDist. Tay Shu Chian for editing the manuscript. Diego Pitta de Araujo for making the illustration diagrams. Ince Tan for providing the ZT-GFP cells.

## Author Contributions

S.K.W. conceived the project with input from Yuting Lou and A.S.Y. C.Z.H. and S.K.W. performed all experiments except traction force microscopy and immunoblotting, which were shared with J.X. and C.B.H. D.L. and D.S.T. provided cancer pathology specimens and performed pathological analysis. F.Q.S. and Yuan Lin developed the theoretical model and conducted simulations and quantitative analyses for fracture dynamics and actomyosin fiber alignment. T.H. and Yuting Lou performed spheroid velocity and jamming transition analyses. P.M. provided the BISM code and assisted with BISM analysis. M.H. Nai, Yuting Lou, and C.T. Lim contributed to AFM-based tension measurements. C.B.H., I.Y., and M.S. performed traction force analysis and 3D StarDist AI–based nuclear segmentation. S.V. and A.S.Y. provided input on cadherin and myosin experiments. S.K.W. wrote the manuscript, with F.Q.S. writing the modelling sections. C.B.H., Yuan Lin, C.Z.H., Yuting Lou, and A.S.Y. edited the manuscript. J.C. contributed to imaging and microscope setup. All analyses were led by S.K.W., with assistance from C.Z.H. All authors provided feedback on the manuscript. S.K.W. supervised the study.

## STAR⍰Methods

### Key resources table

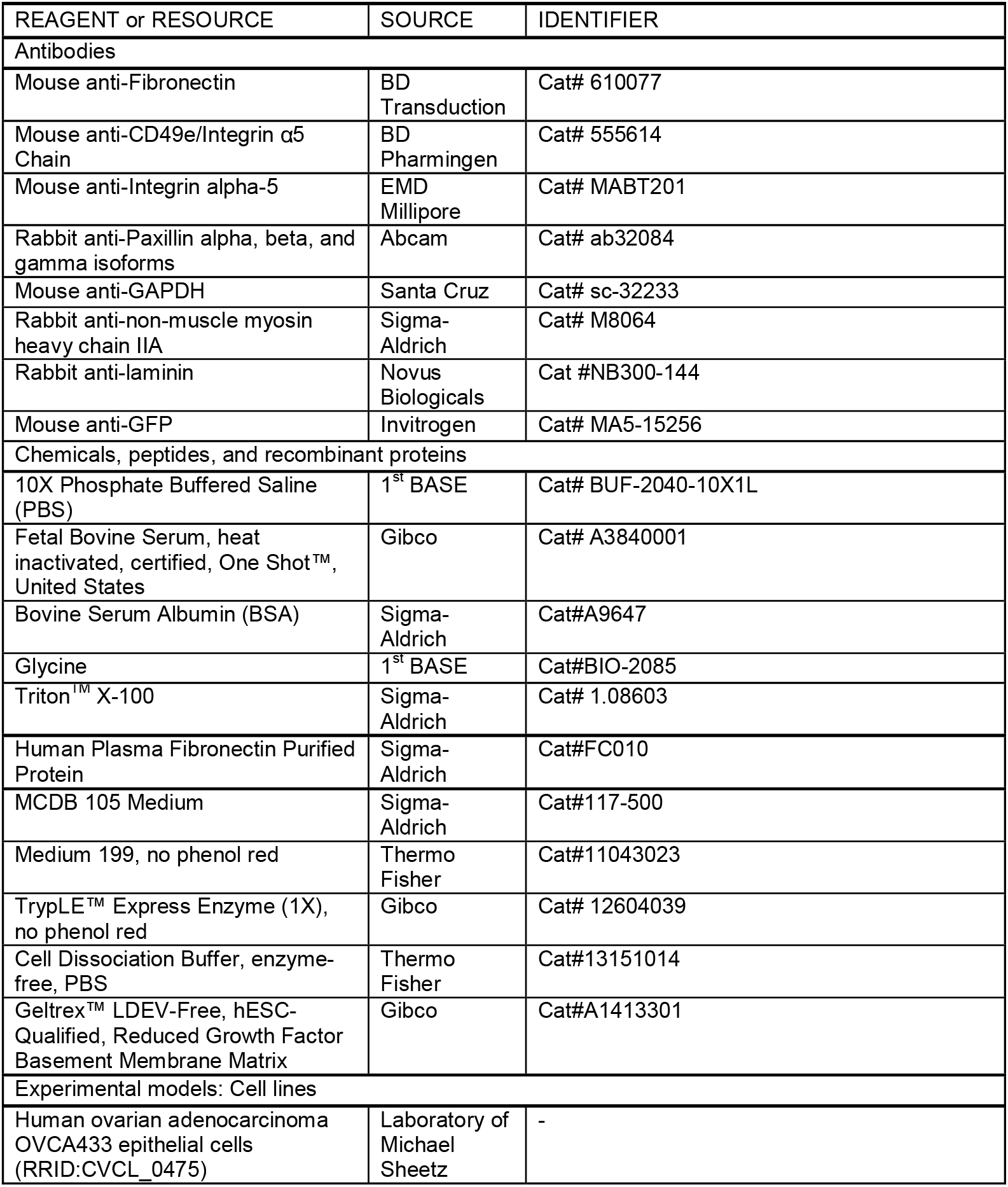
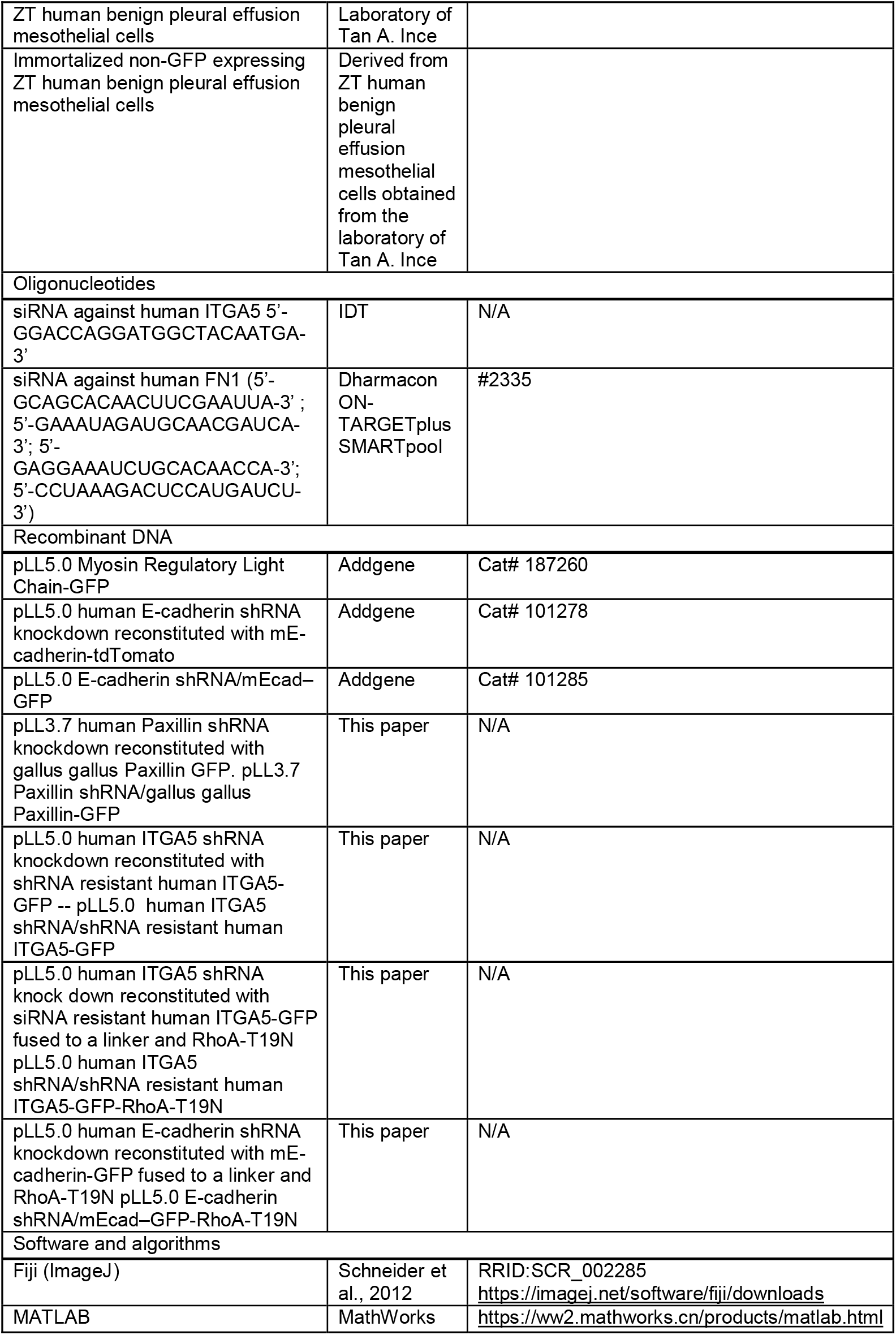
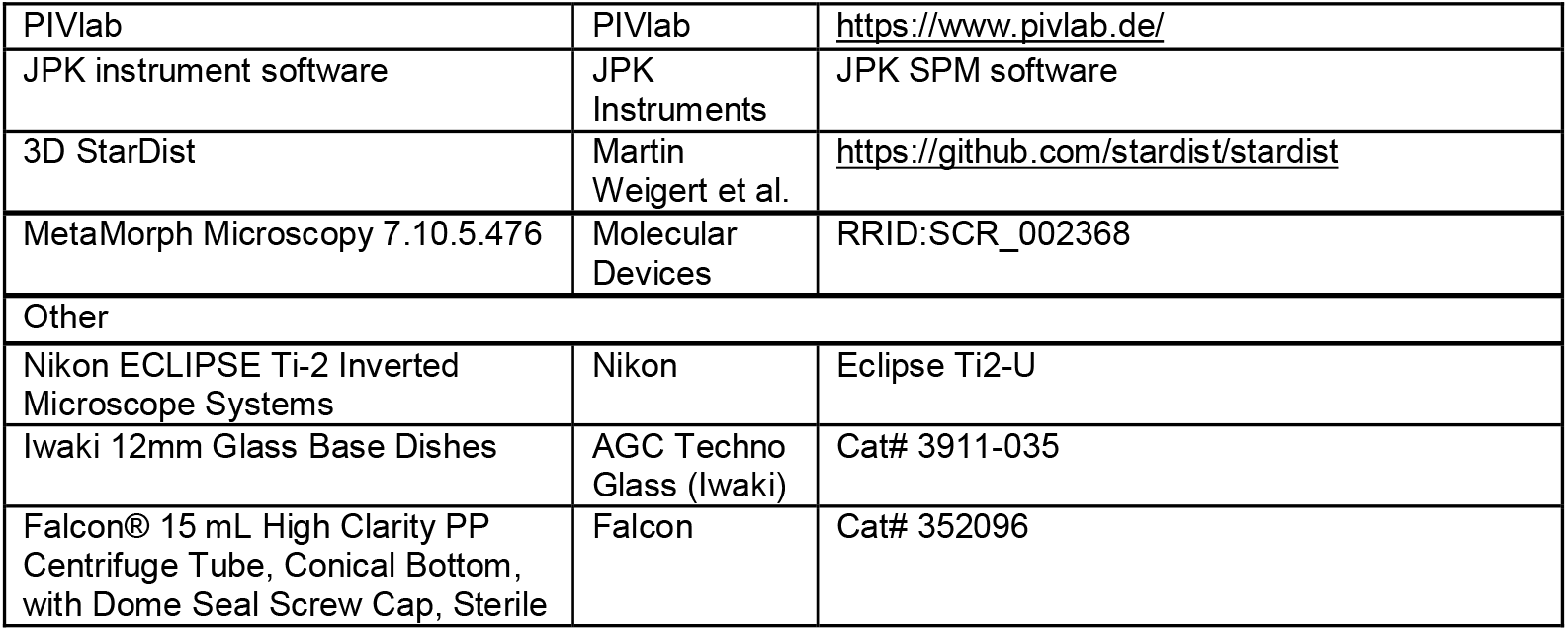

### Experimental models and study participant details

#### Cell culture

OVCA433 ovarian serous adenocarcinoma cell line and the ZT mesothelial cells were cultured in a 1:1 ratio of medium MCDB 105 (Cell Applications Inc.) and Medium 199 (GIBCO) supplemented with 10% FBS (GIBCO). The OVCA433 cell lines were obtained from Michael Sheetz’s laboratory. ZT mesothelial cells were obtained from a benign pleural effusion, a gift from Ince Tan’s laboratory. These cells were immortalized by ectopic expression of SV40 T antigen and overexpression of human telomerase (fused to GFP), as previously described ^13^. Non-GFP expressing ZT cells were obtained by sorting out non-fluorescence ZT cells, and they were subsequently immortalized by transfecting them with SV40 T-antigen expressing plasmid (Addgene). Cells were tested negative for mycoplasma with MycoStrip (Invivogen).

### Human tissue sections

Hematoxylin and eosin (H&E)-staining sections of human ovarian cancer sections were obtained from archived patients in the National University Hospital, Singapore. All samples were obtained with approval by Singapore National Healthcare Group’s Domain Specific Review Board under the reference number 2013/00705. A senior pathologist reviewed the specimens to identify the ovarian tumour invading the peritoneum.

#### Plasmids, siRNA and shRNA

For the expression of fluorescence transgenes we subcloned shRNA-resistant transgene fused with the fluorescence tag into the multiple cloning site of either lentiviral pLL3.7 or pLL5.0 vectors by recombination. The shRNA targeting the corresponding endogenous proteins is cloned into the lentiviral pLL vectors between the XhoI and HpaI restriction sites to prevent overexpression artefacts. The fluorescent transgenes are designed to be shRNA resistant by using a non-human transgene or codon change with site-directed mutagenesis. The lentiviral constructs generated are 1. pLL3.7 human Paxillin shRNA knockdown reconstituted with gallus gallus Paxillin GFP. pLL3.7 Paxillin shRNA/gallus gallus Paxillin-GFP, 2. pLL5.0 human ITGA5 shRNA knockdown reconstituted with shRNA resistant human ITGA5-GFP -- pLL5.0 human ITGA5 shRNA/shRNA resistant human ITGA5-GFP. Additionally, a linker (5′ - GGGTCTACATCTGGATCTGGGAAGCCGGGTTCTGGTGAGGGTTCT-3′) is fused between fluorescence transgenes and the RhoA dominant negative mutants for the following constructs. 3. pLL5.0 human ITGA5 shRNA knock down reconstituted with siRNA resistant human ITGA5-GFP fused to a linker and RhoA-T19N pLL5.0 human ITGA5 shRNA/shRNA resistant human ITGA5-GFP-RhoA-T19N, 4. pLL5.0 human E-cadherin shRNA knockdown reconstituted with mE-cadherin-GFP fused to a linker and RhoA-T19N pLL5.0 E-cadherin shRNA/mEcad–GFP-RhoA-T19N.

The following constructs were published previously. 1. pLL5.0 Myosin Regulatory Light Chain-GFP, 2. pLL5.0 human E-cadherin shRNA knockdown reconstituted with mE-cadherin-tdTomato and 3. pLL5.0 E-cadherin shRNA/mEcad–GFP used was as previously described^19^, 4. Lentiviral H2B-mNeon is a gift from Rong Li’s lab.

Fibronectin 1 and Integrin alpha-5 were silenced by siRNA directed against human ITGA5 5’-GGACCAGGATGGCTACAATGA-3’ and Dharmacon Smartpool siRNA directed against human FN1( 5’-GCAGCACAACUUCGAAUUA-3’; 5’-GAAAUAGAUGCAACGAUCA-3’; 5’-GAGGAAAUCUGCACAACCA-3’; 5’-CCUAAAGACUCCAUGAUCU-3’) respectively.

### Method details

#### Heterotypic spheroid-monolayer coculture

IWAKI 35mm dishes (12mm glass diameter) were coated with 1:100 Human Fibronectin plasma (Sigma-Aldrich) in PBS and incubated at 4°C overnight. For Matrigel conditions, 20µl of Geltrex was added to the glass base and evenly spread. Subsequently, ZT mesothelial cells were seeded and incubated overnight at 37°C to form a confluent monolayer.

Spheroids of OVCA433 were prepared by dissociating cells with trypsin-free Cell Dissociation Buffer, PBS (Gibco), and incubated in suspension in 2ml of 1:1 ratio of medium MCDB 105 (Cell Applications Inc.) and Medium 199 (Gibco) with 10% FBS (Gibco) for 30 mins twice at 37°C to remove trace amounts of cell dissociation buffer and promote spheroid formation. Spheroids of approximately 50µm size were imaged and analysed. Spheroids were transferred to a dish containing a confluent monolayer and incubated for 15 min to allow spheroids to settle before live imaging. Imaging was performed using Nikon W1 Spinning Disk confocal microscope with the integrated Perfect Focus System on a 60x water-immersion magnification objective. Brightfield, GFP, and RFP images were captured every 3 min for 12 h. After live imaging, spheroids that invaded the monolayer were quantified by viewing data obtained using MetaMorph Microscopy Automation and Image Analysis Software. To quantify the mesothelial clearance area, the non-fluorescent area created by the invading spheroid in the GFP mesothelial monolayer images was measured at 24Lh and divided by the initial area of the cancer spheroid 2D size at time 0. All measurements were taken using ImageJ software.

#### Protein coupling of microbeads

PolyLink Protein Coupling Kit (Polysciences, Inc) was used to prepare 50µl of fluorescence beads solution. 200µg of Wheat Germ Agglutinin was added to the activated beads suspension and incubated for 2 hr at room temperature. Subsequently, the beads suspension was sonicated for 15 min and immediately added to a dish containing a confluent monolayer and centrifuged until no floating beads could be observed. Prepared OVCA433 spheroids were transferred to the dish and incubated for 15 min to allow spheroids to settle before live imaging.

Multicellular spheroids of OVCA433 were prepared by dissociating cells with trypsin-free Cell Dissociation Buffer, PBS (Gibco) and incubated in suspension in 2ml of 1:1 ratio of medium MCDB 105 (Cell Applications Inc.) and Medium 199 (Gibco) with 10% FBS (Gibco) for 30 mins twice at 37°C to remove trace amounts of cell dissociation buffer and promote spheroid formation. Spheroids were transferred to a dish containing a confluent monolayer and incubated for 15 min to allow spheroids to settle before live imaging. Imaging was performed using Nikon W1 Spinning Disk confocal microscope with the integrated Perfect Focus System on the 60x water-immersion magnification objective. Brightfield, GFP, and RFP images were captured every 3 min for 12 h. Spheroids that invaded the monolayer were quantified with MetaMorph.

#### Plasmids, siRNA and shRNA

For the expression of fluorescence transgenes we subcloned shRNA-resistant transgene fused with the fluorescence tag into the multiple cloning site of either lentiviral pLL3.7 or pLL5.0 vectors by recombination. The shRNA targeting the corresponding endogenous proteins is cloned into the lentiviral pLL vectors between the XhoI and HpaI restriction sites to prevent overexpression artefacts. The fluorescent transgenes are designed to be shRNA resistant by using a non-human transgene or codon change with site-directed mutagenesis. The lentiviral constructs generated are 1. pLL3.7 human Paxillin shRNA knockdown reconstituted with gallus gallus Paxillin GFP. pLL3.7 Paxillin shRNA/gallus gallus Paxillin-GFP, 2. pLL5.0 human ITGA5 shRNA knockdown reconstituted with shRNA resistant human ITGA5-GFP -- pLL5.0 human ITGA5 shRNA/shRNA resistant human ITGA5-GFP. Additionally, a linker (5′ - GGGTCTACATCTGGATCTGGGAAGCCGGGTTCTGGTGAGGGTTCT-3′) is fused between fluorescence transgenes and the RhoA dominant negative mutants for the following constructs. 3. pLL5.0 human ITGA5 shRNA knock down reconstituted with siRNA resistant human ITGA5-GFP fused to a linker and RhoA-T19N pLL5.0 human ITGA5 shRNA/shRNA resistant human ITGA5-GFP-RhoA-T19N, 4. pLL5.0 human E-cadherin shRNA knockdown reconstituted with mE-cadherin-GFP fused to a linker and RhoA-T19N pLL5.0 E-cadherin shRNA/mEcad–GFP-RhoA-T19N.

The following constructs were published previously. 1. pLL5.0 Myosin Regulatory Light Chain-GFP, 2. pLL5.0 human E-cadherin shRNA knockdown reconstituted with mE-cadherin-tdTomato and 3. pLL5.0 E-cadherin shRNA/mEcad–GFP used was as previously described^19^, 4. Lentiviral H2B-mNeon is a gift from Rong Li’s lab.

Fibronectin 1 and Integrin alpha-5 were silenced by siRNA directed against human ITGA5 5’-GGACCAGGATGGCTACAATGA-3’ and Dharmacon Smartpool siRNA directed against human FN1( 5’-GCAGCACAACUUCGAAUUA-3’; 5’-GAAAUAGAUGCAACGAUCA-3’; 5’-GAGGAAAUCUGCACAACCA-3’; 5’-CCUAAAGACUCCAUGAUCU-3’) respectively.

#### Computational fracture analysis of microbeads

Analysis of microbeads at regions of mesothelial fracture was performed as previously described with minor modifications and additional quantifications^23^. Particle image velocimetry (PIV) analysis (PIVlab package^60^ in MATLAB) was used to quantify the flow fields in the top-viewed cell layer microbeads time-lapse datasets. The PIV analysis was carried out for specific duration segments of these datasets where the field of view was fixed. The time interval of each slice is 30 seconds. We then carry out the PIV analysis by dividing the images into 64×64 pixel interrogation windows with 50% overlap on the first pass, 32×32 pixel interrogation windows with 50% overlap for the second pass, and 16×16 pixel interrogation windows with 50% overlap for the third pass. Next, we select velocity threshold limits for post-processing the resulting velocity vector fields, smooth the data, and use bicubic interpolation for missing vectors. After calibrating the results, velocity magnitude 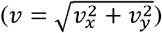, normal strain rate 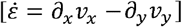^28^ and shear strain rate 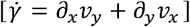 were calculated, where *x* and *y* are the directions of vectors while *v*_*x*_ and *v*_*y*_ represent the corresponding mesothelium velocity components^23^.

#### Traction force microscopy

TFM substrate was prepared as previously described ^61^ with minor modifications. Briefly, PDMS with an average stiffness of 15 kPa was prepared by dispensing 40 µL of 1:1 CyA and CyB mixture on the 14 mm well of the 35 mm glass bottom dish (Cellvis), spin-coating the dish at 1000 rpm for 1 min and letting it cure at 80°C for 2 h. PDMS surface was then silanised with 5% APTES in absolute ethanol, functionalised with filtered and sonicated 0.2% (v/v) 0.2 µm dark red fluorescent carboxylate-modified microspheres (FluoSpheres). Beads were inactivated by 0.1M Tri’s treatment for 10 min. The substrate was coated with 100 µg/mL of poly-D-lysine for 1 hour, followed by the adding 50 µg/ml (1:100) of human plasma fibronectin (Sigma-Aldrich) for overnight incubation at 4°C.

Subsequently, ZT mesothelial cells were seeded and incubated at 37°C to form a confluent monolayer. Multicellular spheroids of OVCA433 were prepared and transferred to the confluent monolayer and incubated for 15 min to allow spheroids to settle before live imaging. Imaging was performed using Nikon W1 Spinning Disk confocal microscope with the integrated Perfect Focus System on the x60 water magnification objectives. After 12 h, 10% SDS was added to remove cells, and the substrate was left alone to recover to the resting state for an hour before acquiring reference images.

#### TFM Analysis

The fluorescent bead images were maximum projected for each timepoint and subsequently contrast-enhanced and registered to the maximum projected reference images using MultiStackReg. PIVlab^62^ add-on for MATLAB was used to calculate the displacement field from the registered bead images using three passes of cross-correlation with the interrogation windows as follows: 128(64), 64(32), and 32(16). Outlier vectors with abnormally large values were removed post-processing. FTTC, an ImageJ plugin for a Fourier transform-based, semi-infinite half-space method, was used to reconstruct the force field from the displacement field^63^. Customised codes were written to generate the force map or superimpose the traction force over the cell image. For downstream analysis requiring quantification for each cell, a code was customised to import ImageJ ROIs into MATLAB to calculate the parameters for each ROI. Radial traction stress is analysed as previously described, with the middle of initially spreading cells or fracture site being the point of reference^64^. Statistical analysis was performed using one-way ANOVA in GraphPad Prism.

#### Bayesian inversion stress microscopy and spheroid surrounding border analysis of tensile stresses

BISM analysis was performed on traction force microscopy data as previously described^26,36^. Next, we identified the spheroid’s outline from brightfield live imaging and created masks for a 10-micron border around spheroids that invaded or did not invade with a MATLAB script. Stress values from the BISM analysis in each 10-micron region bordering the spheroid are obtained to calculate the average stress.

#### Immunofluorescence and microscopy

Cells were fixed at 4 ° C with either −20 ^°^C methanol on ice for 5 min or with 4% paraformaldehyde in cytoskeleton stabilization buffer (10 mM PIPES at pH 6.8, 100 mM KCl, 300 mM sucrose, 2 mM EGTA and 2 mM MgCl2) on ice for 20 min and subsequently permeabilized with 0.2% Triton-X in PBS for 5 min at 4 ^°^C^65-69^.

For structured illumination microscopy, spinning-disc confocal microscopy (Yokogawa) coupled with the Live SR module (GATACA Systems), Nikon Eclipse Ti-E inverted microscope with Perfect Focus System, controlled by MetaMorph software (Molecular device) supplemented with a 100x oil 1.45 NA CFI Plan Apo Lambda oil immersion objective and sCMOS camera (Prime 95B, Photometrics).

For lattice light sheet microscopy, Zeiss Lattice Lightsheet 7 is equipped with a 10x 0.4 NA illumination objective at a 30-angle to cover glass and a 48x 1.0 NA detection objective at a 60-angle to cover glass. Pre-defined Sinc3 beams with 30mm length x 1000nm thickness were used for image acquisition. Images were captured with Pco.edge 4.2 CLHS sCMOS camera and processed with Lattice Lightsheet 7 Processing Module.

#### Antibodies and blocking peptide

Primary antibodies in this study were: mouse monoclonal antibody against human Fibronectin (1:100; BD Transduction Laboratories; catalogue number 610077); mouse monoclonal antibody against human CD49e/Integrin α5 Chain (1:1000 for western blotting; BD Pharmingen; catalogue number 555614); mouse monoclonal antibody against human Integrin alpha-5 (1:100 for immunofluorescence; EMD Millipore; catalogue number MABT201, clone number SNAKA51); rabbit monoclonal antibody [Y113] against mouse and human Paxillin alpha, beta, and gamma isoforms (1:200 for immunofluorescence; Abcam; catalogue number ab32084); mouse monoclonal antibody against GAPDH (1:1000 for western blotting; Santa Cruz Biotechnology; catalogue number sc-32233); rabbit polyclonal antibody against non-muscle myosin heavy chain IIA (1:200 for immunofluorescence; Sigma-Aldrich; catalogue number M8064); rabbit polyclonal antibody against laminin (1:100 for immunofluorescence; Novus Biologicals; catalogue number NB300-144); Mouse monoclonal antibody against native and denatured forms of GFP (1:1000 for western blotting; Invitrogen; catalogue number MA5-15256, clone number GF28R).

#### Generation of micropatterned substrates

Micropatterned substrate was fabricated by stencil patterning. Polydimethylsiloxane (PDMS) (Sylgard 184 kit, Dow Corning) was applied onto a photoresist mould, containing circular micropatterns. A mixture of elastomer to crosslinker in a 10:1 ratio (w/w) was used, followed by curing at 80°C for 2 hours. The resulting crosslinked PDMS layer was carefully detached, and stamps were manually crafted. Inverting the PDMS stamp, it was positioned on an uncoated 35mm μ-dish (ibidi GmbH) with hydrophobic properties. Next, Norland Optical Adhesive 73 (NOA-73, Norland Inc.) was applied along one edge of the stamp. Through capillary action, NOA-73 flowed into the gaps between the PDMS stamp and the dish, effectively sealing the stamp on all sides. Curing the NOA-73 stencil took place under ultraviolet light for 15 seconds. Following removal of the PDMS stamp, the stencil and dish were subjected to incubation with fluorescence fibrinogen mixed fibronectin at 50 μg/ml concentration in PBS. This incubation occurred overnight at 4°C following a brief degassing at 10 mbar. After the incubation, the fibronectin solution was removed, and the stencil was detached. The dish’s printed bottom was treated with 0.2% Pluronic acid-H2O for 30 minutes as a passivation step. Finally, the passivated dishes underwent three washes with PBS before cell seeding.

#### Spheroid nucleus segmentation with StarDist 3D deep learning model and cell tracking by IMARIS

The spheroids’ image stacks underwent manual labelling using the Labkit plugin in ImageJ. These labels were subsequently employed as the ground truth masks for the StarDist 3D segmentation (cite Weigert). A subset of the labelled stacks, selected randomly, was allocated for training (N_*train*_=19) and testing (N_*test*_=2) datasets. To augment the data, various techniques, including random flips, rotations, and intensity modifications, were applied during training to expand the pool of training examples. The initial model parameters were set using the pre-trained StarDist model weights.

Throughout the training process, the last five Conv3D layers of the model were reinitialized and trained over 100 epochs. A learning rate of 10^-4^ was employed, along with a patch size of 8×96×96. All other training parameters were kept consistent as previously described. Following this, the best-performing model, determined by the lowest test dataset loss, underwent further fine-tuning for an additional 10 epochs. During this fine-tuning phase, a reduced learning rate of 10^-6^ was employed. All training and fine-tuning procedures were executed on a shared Jupyter notebook server equipped with an NVIDIA RTX 6000 GPU.

After model training, the trained model was deployed for predictions on new spheroid images. This inference step was carried out using a Google Colab notebook with GPU acceleration (Google, Google Colaboratory. Retrieved from https://colab.google (2025)). After executing the prediction code, the resultant labels were compared with the original image stack to assess segmentation accuracy. Following this evaluation, the autoregressive motion algorithm was applied to enable automated tracking of the segmented H2B mNeon nuclei time-lapse images in IMARIS to generate nuclei tracks. These tracks were then subjected to manual refinement across all frames to accurately capture the movement of spheroid cells.

#### Pair-wise mean square displacement (MSD) analysis of cell nucleus

Given a time *t*, the distance of two nuclei *i* and *j* is denoted as d*x*_*i,j*_ (*t*). The displacement of this distance over an time interval *dt* is d*x*_*i,j*_(*t, dt*)= d*x*_*i,j*_ (*t+dt*)-d*x*_*i,j*_ (*t*). Then, the pair-wise MSD <d*x*^2^>(*dt*) is the squared distance averaged over all time points and all nuclei pairs. MSD increases with *dt* with a power *s* such that <d*x*^2^> ∼ *dt*^*s*^, where *s*(*dt*) is referred to as local slope of MSD power in the main text. A local slope of 1 across all timescale *dt* indicates a random diffusion of particles.

#### Cell migration persistency analysis

Let ***x***_*i*_ be the position of cell nuclei *i* in the XYZ (Cartesian) coordinate space, where the origin of the space ***x***_0_ is the averaged positions of all cells in spheroids. Then, the velocity of nucleus *i* is calculated as the distance of the nucleus between two consecutive frames: ***v***_*i*_(*t*)=***x***_*i*_(*t*+*dt*)-***x***_*i*_(*t*), where *dt* = 3 minutes, the time resolution of tracking. The direction of the velocity is ***v***^***l***^_*i*_ *=****v***_*i*_(*t*)*/*|*v*_*i*_(*t*)|, from which the auto-correlation of velocity directions *C*(*dt*)*= <****v***^*l*^_*i*_ (*t*)***v***^*l*^_*i*_ (*t+dt*)*>* could be obtained. The decay of *C* with the lag time *dt* follows an exponential form as *C ∼* exp(-*dt*/*T*), where *T* is the persistent timescale.

#### Z-stacks of spheroid

Let *S* be the total number of stacks to be sliced, and each stack is enumerated as integer *s* ranging from 1 (top) to *S* (base). We first calculate the cumulative distribution *M* of the cell nuclei in the spheroid along the *Z* direction, then *M*=0 corresponds to the top surface and *M* =1 at the bottom surface. Next, we determine the upper bound *z*_*s*_ and lower bound *z’*_*s*_ of a stack *s* in such a way that *M*(*z*_*s*_)= (*s*-1)/*S* and *M*(*z’*_*s*_)= min(1, (*s*-1)/*S* + *h*), where *h* is a predefined effective thickness (0 < *h* < 1). Note that a fixed *h* imposes a similar number of cells within each stack and excludes the bias from the population number in the statistics. Finally, we select the cells with their vertical position *z* within the range of [*z*_*s*_, *z’*_*s*_] for a stack *s*. In the main text, the total number of stacks *S* is 6, and the effective thickness *h* is 0.3.

#### Interface Energy Formulation

During spheroid wetting and invasion into the cell monolayer, the free energy change (Δ*G*) of the system is due to variations in the interface energies and cell polarization energy, i.e.,

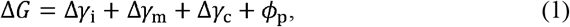

in which Δ*γ*_*i*_, Δ*γ*_*m*_ and Δ*γ*_*C*_ represent changes in the cadherin-mediated spheroid cell-cell, mesothelial cell-cell, and integrin-based spheroid-mesothelium interface energies, respectively. *ϕ*_p_ is the work done by the polarization force in spheroid cells. Physically, these aforementioned interface energies depend on mesothelial cell cortical tension *T*_0_, spheroid cell tension *T*_*C*_, mesothelial adhesion strength *W*_0_, spheroid adhesion strength *W*_*C*_ and integrin-based adhesion energy *W*_*I*_ between spheroid leader cell and mesothelial cells.

During wetting, i.e. when the spheroid leader cell is in contact with the mesothelium, integrin is recruited to the interface (Fig. 3a,b) and increases the adhesion strength *W*_*I*_. Here, such gradual enhancement of adhesion is described by

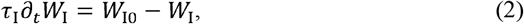

where *τ*_I_ represents the timescale for integrin recruitment, *W*_I0_ is the maximum interface adhesion energy density that can be achieved.

It is widely reported that active contraction of cells could be triggered by mechanical signals such as local stretching and indentation ^46,70^. To incorporate this important feature into our model, a first-order coupling between the cortical tension *T* and the length *L*_in_ of spheroid-mesothelium adhesive interface ^71,72^was adopted as

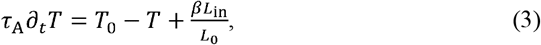

where *T*_0_ is the static mesothelial cortical tension, *L*_0_ is a reference length, *τ*_A_ characterizes the reaction timescale for myosin recruitment and activation, and *β* is the tension provided by the myosin recruitment per unit length of integrin adhesion. Evidently, under a fixed interface length, the maximum mesothelium contractility that can be induced by myosin recruitment is simply 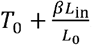.

During wetting, formation or elongation of intercellular junctions would bring in free energy change in the system. Since active constriction is only found at the apical side of the mesothelium, the interface energy changes are Δ*γ*_m_ = −*W*_0_ *δL*_m_, Δ*γ*_*C*_ = −*W*_C_ *δL*_C_ and

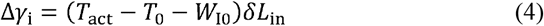

where *T*_aCt_ is the increased mesothelium apical tension in the equilibrium state. Considering the steady-state solution of Eq. (3), the apical tension change is simply 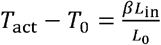. At the same time, as spheroid cells move towards the mesothelium, free energy change induced by polarization work is *ϕ*_p_ = −*F*_p_ *δL*_m_.

At steady-state, the change of system free energy according to Eqs. (1-4) is

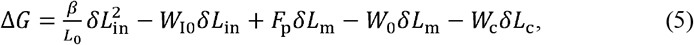

where *δL*_in_, *δL*_m_ and *δL*_C_ are the length change of spheroid-mesothelial integrin junction, mesothelial E-cad junction and spheroid E-cad junction, respectively, *F*_p_ is the polarization force in spheroid cells (Supplementary Fig. 3b). For the spheroid itself, the free energy change reduces to Δ*G*_C_ = −*W*_I0_ *δL*_in_ - W_C_ *δL*_C_, which shows a competition between the integrin interface energy and spheroid E-cad interface energy. This implies that the shortening of spheroid E-cad junction due to the favoring of spheroid-mesothelial junctions lengthening, promotes the wetting of spheroid on the mesothelium^73^. Indeed, experiments with spheroids sorted for higher E-cadherin expression, which prevents cell-cell contact shrinkage^74^and reduces spheroid invasion (Fig. 4j,k, Supplementary Fig. 3k,l). In the scenario of single-cell wetting, we have *δL*_C_ = 0. Under such circumstances, the mesothelial free energy change is 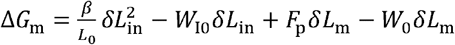, indicating the fracture of mesothelial E-cad junction (represented by the vanishing mesothelial junction length due to *γL*_m_) is regulated by mesothelial apical constriction, spheroid-mesothelium adhesion, and spheroid cell polarization. Specifically, considering the complete rupturing condition *δL*_m_ ≤ -*L*_m0_ (with *L*_m0_ being the initial length of the mesothelial junction), the critical condition for spontaneous mesothelium fracturing (Fig. 5a) is

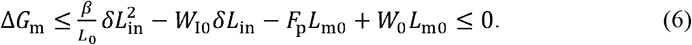

For this equation to have a solution, we must have

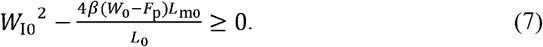

Then, the critical condition for mesothelium fracture (Fig. 5a) can be obtained as

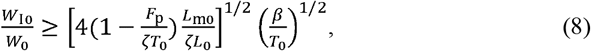

where 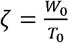 defines the initial mesothelial tissue state.

#### Active Foam Formulation

In our model, cell movement and deformation are captured by the position change of a set of vertices, describing the shape of each cell as

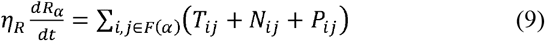

where *R*_*α*_ is the position vector of vertex *α,t* represents the time, _*η*__R_ is the friction coefficient associated with vertex movement, *F* (*α*) represents the set of junctions sharing vertex *α,N*_*ij*_ is the normal force (due to osmotic pressure ^54^) acting on the junction *ij, T*_*ij*_ is the active cortical tension along the tangential direction of junction *ij* (Supplementary Fig. 3b). *P*_*ij*_ is the polarization force within the spheroid leader cell. Note that the effective tension on adherent junctions is affected by the strength of cell-cell adhesion. For example, the magnitude of effective junctional tension *ij* between cell *a* and cell *b* is expressed as 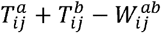, where 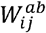 is the adhesion strength on segment *ij* between two cells. The asymmetry of surface forces induced by cell adhesion elongates the interface and drives the spreading of the spheroid leader cell on the mesothelium. The physical meaning and adopted value of different parameters are summarized in Table S1.

#### Initial Configuration

Four spheroid cells that formed a cell cluster and were designated near a mesothelium cell layer were used to study the dynamic invasion process. Periodic boundary conditions are applied to the left and right sides of the mesothelium monolayer during the simulation. As the cancer cells are supposed to form a spheroid, a relatively high adhesion energy among themselves is adopted (Table S1).

On the other hand, a very weak interface adhesion between cancer spheroid and mesothelial cells (i.e., W_I0_⁄*T*_0_ = 0.05) is applied at the beginning of the simulation, given that stable spheroid-mesothelial attachment takes time to form (after their initial contact). A slightly lower E-cadherin adhesion energy (i.e., 0.8*W*_0_) between mesothelial cells is also adopted. Before the actual simulation, the system was allowed to relax the steady state.

Once the simulation starts, a dynamic spheroid-mesothelium adhesion strength (given by Eq. (3)) is invoked. Driven by the increased spheroid-mesothelial adhesion, the spheroid leader cell spreads more on the mesothelium, triggering its apical contraction and junction rupture and eventually leading to spheroid invasion into the mesothelial layer.

Based on our experimental observations, *L*_0_ and *T*_0_ are taken to be 20 mm and 5 nN, respectively. According to previous studies ^72,75^, the friction coefficient *η* _R_ could vary from 0.01 to 0.1 N·s·m^-1^; here, we take a high friction coefficient (0.1 N·s·m^-1^) for mesothelial cells as they are attached to the substrate and a low friction coefficient (0.05 N·s·m^-1^) for the floating spheroid cells. Since the integrin adhesion timescale is believed to be significantly shorter than that for apical constriction ^72,76,77^, we choose *τ*_I_ = 20 s and *τ*_A_ = 100 s in the simulation. The values of all dimensionless parameters are listed in Table S1.

**Table S1.** Description of dominating parameters.

| Parameter | Physical description | Range | Data Estimation |
| --- | --- | --- | --- |
| $\frac{L_c}{L_0}$ | Characteristic spheroid size | 1 | Measurement |
| $\frac{T_c}{T_0}$ | Spheroid cortical tension | 0.5 | Measurement |
| $\frac{\beta}{T_0}$ | Integrin-based apical tension strengthening | 0.5~5 | Literature<sup>71,72</sup> |
| $\frac{W_0}{T_0}$ | Mesothelial adhesion strength | 1.2 | Shape estimation |
| $\frac{W_c}{T_c}$ | Spheroid adhesion strength | 1.1 | Shape estimation |
| $\frac{P_0 L_0}{T_0}$ | Static osmotic pressure | 8000 | Literature<sup>78</sup> |
| $\frac{F_p}{T_0}$ | Cell polarization force | 0.72 | Literature<sup>79</sup> |

#### Onset of Tissue Fracture

At the early stage of spheroid cell invasion, the mesothelial junction below spheroid cells is shortened by activated apical contraction. In our simulation, we set a mesothelial junction to be completely ruptured when its length is below a small threshold value (i.e., ≤ *L*_0_⁄100), corresponding to the onset of tissue fracture in our experiments. In this situation, the remaining central vertices are copied and divided equally between the cells on both sides.

#### Vertex Motion and Contact Formation

Movements of vertices in our model are governed by Eq. (1). To maintain the accuracy of the simulation; automatic re-meshing is adopted where a new vertex is added to the middle of the boundary segment when its deformed length is above a critical value (i.e., *L*_0_⁄20).

To capture the growth of adhesive contact between cells in our simulation, vertices in front of each contact tip are checked at every time step. Once a segment from one cell is close enough to the other cell boundary, the two segments (as well as vertices) are combined. In this way, elongation of the contacting spheroid-mesothelial interface is realized.

The shape index of the cell is calculated as 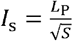 where *L*_p_ is the perimeter of the cell and *s* is the area.

#### Cell stiffness and surface tension measurement by atomic force microscopy (AFM)

The stiffness of the mesothelial cells and spheroids was measured using a Nanowizard IV BioAFM system (JPK Instruments, Germany). Confluent mesothelial cell monolayers were cultured on a glass bottom petri dish while the spheroids were adhered to a poly-d-lysine (PDL) coated glass bottom petri dish. Indentations for surface tension measurements were performed on randomly selected cells with a silicon nitride pyramidal tip (MLCT-D, Bruker; nominal cantilever spring constant k = 0.03 N/m; nominal tip radius 20 nm) using a force of 3 nN at a speed of 5 µm/s. The force curves from more than nine mesothelial cells and multiple cells from nine ovarian cancer spheroids were analysed to evaluate the surface tension. The calculation of cell surface tension was performed as recently described^80-83^. In brief, the cell membrane is assumed to be a thin tensile shell. With an AFM indentation range much larger than cell membrane thickness, the formula used is T = K/2pi, where K is fitted from the linear coefficient of force vs. indentation ranging from 50nm to 300 nm.

### Quantification and statistical analysis

#### Image Processing and Analysis

The images presented were processed with ImageJ (http://rsb.info.nih.gov/ij/). For quantification of apical fluorescent density and line-scan analysis, the mean intensity of the selected apical region was measured from the microscopy images. Line-scan analysis was performed according to the published protocol^67^: A line of standardized length was drawn perpendicular to and centered on cell-cell contacts, and fluorescence intensity of all pixels along the drawn line was determined using the PlotProfile feature on ImageJ. The resulting data was imported into Excel for average calculation. For statistical analysis, raw fluorescence intensity data was used in Prism to analyze co-localization between E-cad GFP spheroid and E-cad tdT mesothelium.For quantification of the phosphoMLC intensity at E-cadherin cell-cell contacts and apical integrin adhesions, segmented lines were drawn on immunostained E-cadherin cell-cell contacts (line width 20) and apical integrin adhesions to obtain multiple regions of interest with FIJI. The corresponding phosphoMLC intensities were measured at these regions of interest.

For quantification of apical mesothelial paxillin intensity, an image mask was created by subtracting the background intensity with the rolling ball radius set to 50 pixels with FIJI, followed by the Gaussian blur with a sigma of 2. Particles with an area larger than 0.1 μm^2^ were considered and segmented for the final mask to measure Paxillin intensity. The area of each focal adhesion was recorded as a region of interest (ROI) in ImageJ, and the mean intensity of the focal adhesions was calculated by subtracting the background intensity obtained from an area of the image without paxillin. For quantifying mesothelial apical F-actin orientation, we measured the alignment of F-actin bundles in cell fluorescence images at 10-degree intervals, using the Directionality Function in Fiji (v. 1.54p).

For quantification of the abundance of fibronectin fibrils at the apical regions of the mesothelium, an intensity threshold mask was created to outline the fibronectin fibrils with FIJI. Their intensity multiplied by the area of these fibronectin fibrils are then compared between the different experimental groups.

For quantification of cell-cell contact crazing, time-lapse image stacks of mesothelial monolayers were analysed using Metamorph with Z-projection. Mesothelial monolayers were first visually verified to be intact before tearing. Timeframes showing the initial cell-cell contact breakdown were determined after playing through each movie and visually assessed for multiple fibrillar bridges and microvoids indicative of crazing.

## Statistical analysis

All data are represented as mean ± SD, unless otherwise stated. Each experiment was independently performed at least 3 times. All statistical analyses were performed in GraphPad Prism. Student’s t test was used to compare datasets with 2 groups; depending on the number of variables, one-way or two-way ANOVA was used to compare 3 or more groups.

## Supplemental information

### Supplementary videos

Video 1. Formation of fibrillar bridges and microvoids at mesothelial E-cadherin (hot orange) cell-cell contacts during spheroid invasion, related to Figure 1.

Video 2. Formation of fibrillar bridges (E-cad tdTomato, red) and microvoids at mesothelium (GFP, green) grown on matrigel during spheroid invasion, related to Figure 1.

Video 3. Apical constriction of a mesothelial cell expressing myosin regulatory light chain (MRLC-GFP, green) and E-cadherin tdTomato (red) during the spheroid invasion, related to Figure 2.

Video 4. Mesothelial apical fibronectin GFP and mesothelium (E-cad tdTomato, red) undergo fracture induced by an invading spheroid, related to Figure 3.

Video 5. Representative multipanel videos of spheroid invasion of the mesothelium in the three following experimental conditions, related to Figure 3.

1) Fibronectin-1 and 2) α5 integrin siRNA mediated knockdown in mesothelium compared to controls. 3) Mesothelium expressing α5 integrin GFP fused with RhoA-T19N compared to control mesothelium expressing wild-type α5 integrin GFP. Arrows indicate four ends of the mesothelial fracture propagation.

Video 6. Representative multipanel videos of spheroid invasion of the mesothelium in the three following experimental conditions, related to Figure 3.

1) Control mesothelium expressing wild-type α5 integrin GFP, 2) mesothelium expressing α5 integrin GFP fused with RhoA-T19N, and 3) mesothelium expressing α5 integrin GFP fused with RhoA-T19N with addition of exogenous FN1. Arrows indicate four ends of the mesothelial fracture propagation.

Video 7. Representative multipanel video of spheroid expressing H2B-mNeon invading E-cad tdT expressing mesothelium and the corresponding XZ and RZ velocity maps on the right, related to Figure 4.

Video 8. Computational vertex results of spheroid cell invasion and mesothelial fracture with medium intercellular integrin adhesion, related to Figures 5 and 6.

Top left: (*W*_I0_⁄*W*_0_ = 0.9), top right: (*W*_I0_⁄*W*_0_= 0.3), bottom left: (*W*_I0_⁄*W*_0_= 1.5), bottom right: (*W*_0_⁄*T*_0_= 0.6).

